# Targeted repression of DNA topoisomerase I by CRISPRi reveals a critical function for it in the *Chlamydia trachomatis* developmental cycle

**DOI:** 10.1101/2023.03.14.532001

**Authors:** Li Shen, Leiqiong Gao, Abigail R. Swoboda, Scot P. Ouellette

## Abstract

*Chlamydia trachomatis* is an obligate intracellular bacterium that is responsible for the most prevalent bacterial sexually transmitted infections. Changes in DNA topology in this pathogen have been linked to its pathogenicity-associated developmental cycle. Here, evidence is provided that the balanced activity of DNA topoisomerases (Topos) contributes to *Chlamydia* developmental processes. Utilizing catalytically inactivated Cas12 (dCas12) based-clustered regularly interspaced short palindromic repeats interference (CRISPRi) technology, we demonstrate targeted knockdown of chromosomal *topA* transcription in *C. trachomatis* without detected toxicity of dCas12. Repression of *topA* impaired the growth of *C. trachomatis* mostly through disruption of its differentiation from a replicative form to an infectious form. Consistent with this, expression of late developmental genes of *C. trachomatis* was downregulated while early genes maintained their expression. Importantly, the growth defect associated with *topA* knockdown was rescued by overexpressing *topA* at an appropriate degree and time, directly linking the growth patterns to the levels of *topA* expression. Interestingly, *topA* knockdown had pleiotropic effects on DNA gyrase expression, indicating a potential compensatory mechanism for survival to offset TopA deficiency. *C. trachomatis* with *topA* knocked down displayed hypersensitivity to moxifloxacin that targets DNA gyrase in comparison with the wild type. These data underscore the requirement of integrated topoisomerase actions to support the essential development and transcriptional processes of *C. trachomatis*.

**Importance:** We used genetic and chemical tools to demonstrate the relationship of topoisomerase activities and their obligatory role for the chlamydial developmental cycle. Successfully targeting the essential gene *topA* with a CRISPRi approach, using dCas12, in *C. trachomatis* indicates that this method will facilitate the characterization of the essential genome. These findings have an important impact on our understanding of the mechanism(s) by which well-balanced topoisomerase activities enable *C. trachomatis* to adapt to unfavorable growth conditions imposed by antibiotics.

## Introduction

A group of enzymes, namely DNA topoisomerases (Topos), act to correct the altered DNA topology that occurs during DNA replication, transcription, and recombination by causing temporary breaks on the DNA helix to prevent excessive supercoiling that is deleterious (1, 2). Most pathogenic bacteria encode two classes of Topos: (i) type IA (e.g. TopoI or TopA) that cleaves and rejoins single-strand DNA independently of ATP, and (ii) type II (e.g. DNA gyrase and Topo IV) that exerts its effects through ATP-dependent double-strand cleavage. An accepted model in *Escherichia coli* is that the concerted action of these Topos dictate the topological properties of DNA (3-5). Whereas active gyrase holoenzyme (composed of two GyrA and two GyrB subunits) negatively supercoils, monomeric TopA removes excessive negative supercoils and works along with gyrase to control the superhelical density of the chromosome. The active Topo IV holoenzyme (composed of two ParC and two ParE subunits) disentangles replicated DNA and enables segregation of daughter chromosomes. Because Topos are ubiquitous, they are considered to be essential for bacterial viability. Quinolone antibiotics, like moxifloxacin, target gyrase and TopoIV in many bacteria and are widely prescribed to treat serious infections associated with *Enterobacterales, Mycobacterium tuberculosis, Pseudomonas aeruginosa, Moraxella catarrhalis, Chlamydia* species, *Mycoplasma* species, and *Staphylococci* species (6-8). However, emerging mutations in genes encoding gyrase or TopoIV have conferred moxifloxacin resistance during the last decades. Characterizing the function of Topos in, and their effects on, bacterial physiology may facilitate the development of new antibacterial therapies.

*Chlamydia trachomatis* is a Gram-negative bacterial parasite that is the leading cause of bacterial sexually transmitted infections worldwide (9). *C. trachomatis* primarily infects human mucosal epithelial cells, where it grows in a membrane-bound vacuole (named as an inclusion) and exists as functionally and structurally distinct forms (10, 11). These forms mainly include (i) the noninfectious, replicative reticulate body (RB) that has a dispersed chromatin structure, and (ii) the infectious, non-replicative elementary body (EB) that is typified by DNA condensation. EB differentiation to RB is detected by 2 hours after infection (h pi), and this is followed by rapid RB multiplication via an asymmetric polarized division mechanism starting at approximately 10 h pi (12-14). RBs begin to asynchronously undergo secondary differentiation into EBs starting at ∼16 h pi, depending on the serovar. However, when stressed, *Chlamydia* can enter an aberrant growth mode called persistence. Signals that trigger the variations of *C. trachomatis* development remain unknown, but one striking change is the DNA supercoiling (11, 15, 16). The superhelical density of the plasmid peaks at ∼24 h pi and is much higher than that at the early or late developmental stages of *C. trachomatis*. These findings raise the questions of how DNA topology is regulated and what the consequences of topological changes are for the chlamydial developmental cycle.

The genome of *C. trachomatis* comprises a chromosome of ∼1.0 Mbp and a plasmid of 7.5 kbp (17). It has been proposed that chlamydial DNA topology is managed by three Topos (gyrase, TopoIV, and TopA) and certain DNA binding proteins (11, 15, 16, 18). The Topo encoded genes are located in three separate operons on the chlamydial chromosome and are commonly transcribed by RNA polymerase containing the major sigma factor σ^66^ (17, 19). However, they are expressed in temporal fashion through a not-yet-identified mechanism(s). *In vitro*, individual recombinant Topo enzymes could modify the superhelical density of plasmid DNA and affect transcription from selected promoters using the plasmid DNA as templates (16, 18, 19). *In vivo* in *C. trachomatis*, one means of defining DNA supercoiling’s involvement in gene regulation was the use of aminocoumarin (i.e. novobiocin) to relax DNA; the resultant effects on transcription of a gene of interest were then measured using reverse transcription quantitative PCR (RT-qPCR) (19). All these studies, while implicitly acknowledging developmentally regulated changes in DNA topology and Topo expression, did not adequately address the question regarding how TopA in conjunction with type II Topos influences the developmental cycle of *C. trachomatis*. The lack, until recently, of genetic tools is the main cause of such knowledge gaps.

The purposes of the current study were to (i) determine the role of TopA in the chlamydial developmental cycle *in vivo* using CRISPRi for targeted knockdown of *topA*, and (ii) investigate the effects of DNA relaxation on chlamydial growth by overexpressing TopA or using moxifloxacin, a pharmacological gyrase/TopoIV inhibitor. The design allows for investigation of how retention of or interference with TopA function alone or in combination with inhibition of gyrase and TopoIV affects key developmental events in *C. trachomatis*. Through growth and morphology measurements, this work describes the first detailed phenotype of TopA deficiency and indicates the importance of carefully balanced Topo activity for the chlamydial adaptive response to supercoiling levels. It further establishes the utility of CRISPRi in understanding essential gene function in this important pathogen.

## Results

### Chromosomal *topA* can be targeted using CRISPRi

Recently, CRISPRi has been used for targeted gene inhibition in *C. trachomatis* (20, 21). To elucidate the role of TopA, we investigated whether it was possible to target chromosomal *topA* using CRISPRi. We created a new spectinomycin-resistance encoding vector, pBOMBL12CRia(topA)::L2 that uses a modified pBOMB4-Tet-mCherry backbone (22). The pBOMBL12CRia(topA)::L2 contains (i) a tetracycline promoter (P_*tet*_)/repressor controlled dCas12 gene with a weakened ribosome binding site, (ii) a specific *topA*-targeting crRNA sequence controlled by a weakened, constitutive chlamydial *dnaK* promoter, (iii) a *Neisseria meningitis* promoter (P_*Nmen*_)-linked *gfp* gene (23), and (iv) a spectinomycin/ streptomycin resistance gene, *aadA*, to facilitate selection for *C. trachomatis* transformants. The design permits that a specific crRNA directs aTC-inducible dCas12 to a specific DNA target (here, the promoter region of *topA* on the *C. trachomatis* chromosome), where it represses transcription (Fig. 1a). The control vector, designated as pBOMBL12CRia(NT)::L2 contained the same components excepting *topA*-specific crRNA, which was replaced with a scrambled sequence with no homology to any chlamydial sequence. Each vector was transformed into *C. trachomatis* serovar L2 -pL2 (20, 24), resulting in strains L2/*topA*-kd and L2/Nt; both were then used individually to infect HeLa cells.

**Figure 1.**
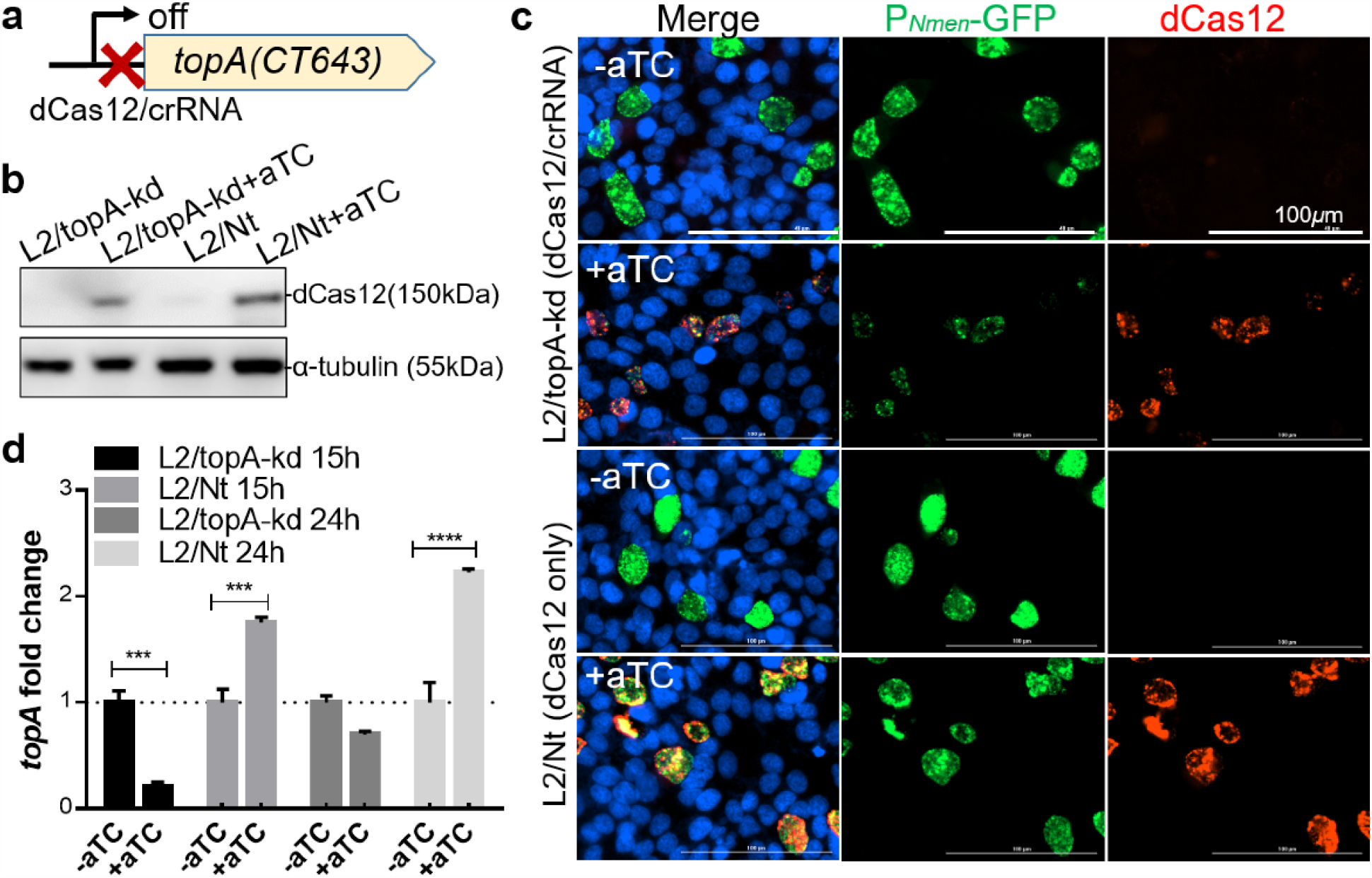
Conditional repression of *topA* transcription in *C. trachomatis*. (**a**) Schematic representation of the strategy used to make a targeted *topA* knockdown through dCas12 and a specific crRNA, whose targeting site is indicated by the red X. (**b**) Immunoblotting analysis of dCas12 expression. *C. trachomatis* L2/topA-kd and L2/Nt (control) infected cells were cultured in medium containing aTC (10ng/mL) for 20 hrs starting at 4 h pi and sampled for immunoblotting with rabbit anti-dCas12 antibody. Host cell α-tubulin was probed with a mouse anti-tubulin antibody and used as a protein loading control. (**c**) Immunofluorescence micrograph of *C. trachomatis* grown in the absence (-aTC) or presence of aTC (+aTC). Fixed cells at 40 h pi were immunolabeled with rabbit anti-dCas12 antibody and visualized with Alexa Fluor 568-conjugated goat anti-rabbit IgG. *C. trachomatis* expressing GFP (green) and dCas12 (red) are shown. Host cell and bacterial DNA were counterstained with 4′,6-diamidino-2-phenylindole (DAPI) (blue). Scale bar=100μm. (**d**) Fold change in relative *topA* transcript levels in the absence or presence of aTC. RT-qPCR was performed with *C. trachomatis* infected cells grown under dCas12-inducing or mock inducing conditions for 11 hrs (to 15h pi) and 20 hrs (to 24h pi) starting from 4 h pi. Chlamydial genomic DNA (gDNA) copy from respective culture was determined by qPCR using primers specific to housekeeping *tufA* gene. Relative quantitation of *topA* specific transcripts were normalized to the gDNA value. The data are presented as the ratio of relative *topA* transcript in the presence of aTC to that in the absence of aTC, which is set at 1 as shown by a black dashed line. The data and standard deviation (SD) of three independent biological replicates are shown. \*\*\**p* < 0.005, \*\*\*\**p* < 0.0001. Statistical significance in all panels was determined by one-way ANOVA followed by Tukey’s post-hoc test.

We first confirmed that dCas12 expression was induced by addition of aTC. *C. trachomatis* infected HeLa cells were cultured in medium without or with aTC (at the concentration of 10 ng/mL or ∼5nM) for 40 hrs. The small amount of aTC used did not have a significant inhibitory effect on the growth of *C. trachomatis* (20, 22, 25). Expression of dCas12 was examined by indirect immunofluorescence assay (IFA) in single cells and by immunoblotting analysis with the lysates of the cell population. We detected the induction of dCas12 expression after adding aTC in both L2/*topA*-kd and L2/Nt cultures (Fig. 1b-c). Most GFP expressing organisms were co-localized with dCas12 signal. These results indicate that dCas12 is inducible and stably present in *C. trachomatis*.

We next determined whether targeted *topA* knockdown occurred in L2/topA-kd using RT-qPCR. Because *topA* is expressed preferentially at the mid-stage, with transcript levels peaking at 14-16 h pi as described previously (19, 26), we expected that aTC addition at 4 h pi would have maximal efficiency in *topA* repression induced by CRISPRi. Indeed, the addition of aTC decreased *topA* transcripts by more than ∼80% and ∼30% at 15 h pi and 24 h pi, respectively, in *C. trachomatis* L2/topA-kd. In contrast, *topA* transcripts were increased approximately two-fold in strain L2/Nt at 24 h pi after adding aTC. The biological relevance of this increase is unknown but may relate to a slight delay in developmental cycle progression when overexpressing the dCas12 (27).

Together, these data demonstrate that *topA* transcription in *C. trachomatis* can be conditionally repressed using CRISPRi. Both *topA*-specific crRNA and the inducible dCas12 expression are necessary and sufficient for successful *topA* knockdown.

### TopA activity is critical for *C. trachomatis* growth

We noted that, under dCas12-inducing conditions, the levels of dCas12 expression and the GFP signal in strain L2/topA-kd were qualitatively lower than those in L2/Nt (Fig. 1). This could have two, non-mutually exclusive, interpretations. First, CRISPRi-induced *topA* repression interferes with *Chlamydia* growth and, second, reduced TopA expression may interfere with P_*Nmen*_-GFP expression from the plasmid. To test these two possibilities, we first generated a series of growth curves to examine chlamydial growth kinetics in the absence or presence of aTC. Growth curves were created by enumeration of the inclusion forming units (IFUs) (equivalent to infectious EB yield) at different time points along the 48 h experimental period after passaging onto a fresh cell monolayer. In the absence of aTC, there was no significant difference in the EB yields between the L2/top-kd and L2/Nt at 24h pi and thereafter (Fig. 2a). Addition of aTC to L2/topA-kd culture resulted in decreased EB yields, which were ∼1/2 log and ∼l log less than those from cultures lacking aTC at 24 and 30 h, respectively. The EB amounts remained ∼2-log lower at 48h pi, indicating an incomplete developmental cycle. Despite the plentiful dCas12 induction in L2/Nt, EBs accumulated at levels similar to (at 24 and 48 h pi, *p*>0.05) or slightly higher (at 30 h pi, *p<0*.*05*) than that of the uninduced conditions. Thus, dCas12 induction alone has a minimal impact on chlamydial growth in the conditions tested. Rather, it is the combination of inducible dCas12 and the targeted crRNA to mediate *topA* repression that causes the growth defect of *C. trachomatis*.

**Figure 2.**
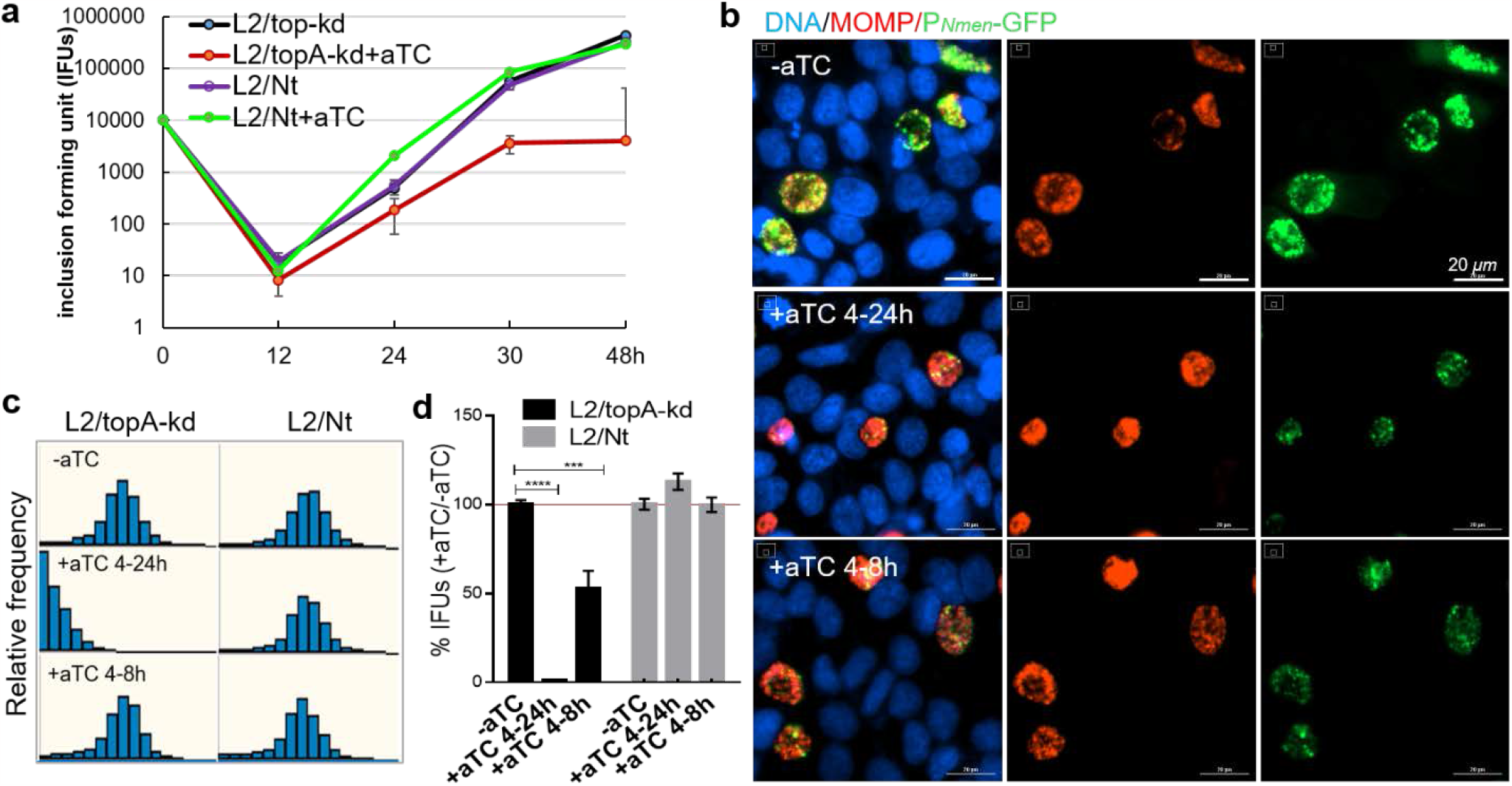
Targeted knockdown of *topA* causes intracellular growth arrest of *C. trachomatis*. (**a**) One-step growth curve of *C. trachomatis*. HeLa cells were infected with *C. trachomatis* L2/topA-kd or L2/Nt at the dose that resulted in 40% cell infection (multiplicity of infection, MOI= 0.4) and cultured in the absence or presence of aTC (at 10ng/mL). Cells sampled at 0, 12, 24, 30, or 48h pi (x-axis) were used for determination of inclusion forming unit (IFUs; y-axis) on fresh HeLa monolayers. IFU values are expressed as the mean ± standard deviation (SD) from triplicate samples. Experiment was repeated three times. (**b**) Representative immunofluorescence images of *C. trachomatis* L2/topA-kd. Infected HeLa cells were grown under the conditions of dCas12 induction for 20 h (+aTC4-24h), transient induction from 4 to 8 h pi (+aTC 4h/-8h), or mock induction (-aTC). Fixed cells at 24 h pi were immunolabeled with monoclonal antibody to *C. trachomatis* major outer membrane protein (MOMP) and visualized with Alexa Fluor 568-conjugated goat anti-mouse IgG. The DAPI-stained DNA (blue), MOMP (red), and *C. trachomatis* expressing GFP (green) are shown. The automated images were obtained with a 20× objective using Cytation 1. Scale bar=20 μm. (**c**) Histogram displays frequency of the individual *C. trachomatis* inclusion sizes that were calculated using Gen 5 software. Graph shows measurement of one representative well with 9 different fields per condition. Three independent trials were performed. (**d**) Relative IFUs in *C. trachomatis* in the absence or presence of aTC for 20 hrs or 4 hrs. Triplicate results in a representative experiment are shown as mean ± SD. Values are presented as the percentage of IFU from dCas12 induced sample to that from respective mock induction sample, which is set at 100 as indicated by a red line. At least four independent experiments were performed. Statistical significance in all panels was determined by one-way ANOVA followed by Tukey’s post-hoc test. \*\*\*\**P*<0.0001; \*\*\**P*<0.001.

*C. trachomatis* grows within the intracellular inclusion niche, whose expansion mirrors the pathogen-host interactions. We sought to examine whether the duration of *topA* knockdown affected the inclusion morphology. Two different culture conditions were chosen: dCas12 induction for 20 hrs (from 4 to 24 h pi to cover the early- and mid-stages) or only 4 hrs (from 4 to 8 h pi to cover the early stage prior to EB accumulation). The size of chlamydial inclusions was measured; in parallel, EB yield was assessed. When dCas12 was induced for 20 hrs (+aTC 4-24h), L2/topA-kd formed smaller inclusions, unlike the control L2/Nt that displayed “normal” large inclusions (Fig. 2b-2c). Consistent with the smaller inclusion size, progeny EBs were decreased by ∼90% in the L2/topA-kd strain (Fig. 2d). Induction of dCas12 for 4 hrs (+aTC 4-8h) had little effect on the inclusion sizes but decreased EB yield by ∼50%, indicating that even a short window of *topA* knockdown can have a measurable effect on chlamydial growth. None of these observed changes were evident for strain L2/Nt. These data suggest that *C. trachomatis* is sensitive to either transient or prolonged *topA* repression induced by CRISPRi, highlighting the critical role of TopA activity in supporting the intracellular developmental cycle of *Chlamydia*.

### P_*Nmen*_-GFP expression is reduced upon *topA* knockdown

As noted above, we observed a weakened GFP signal in *C. trachomatis* L2/topA-kd upon *topA* knockdown, suggesting that the activity of P_*Nmen*_ was potentially inhibited by *topA* repression. Using RT-qPCR, we measured a decrease in *gfp* transcripts in L2/topA-kd, but not in L2/Nt, under dCas12-inducing conditions (Fig. 3a). Thus, changes in P_*Nmen*_-*gfp* levels represents another measurable impact mediated by CRISPRi-induced *topA* repression.

**Figure 3.**
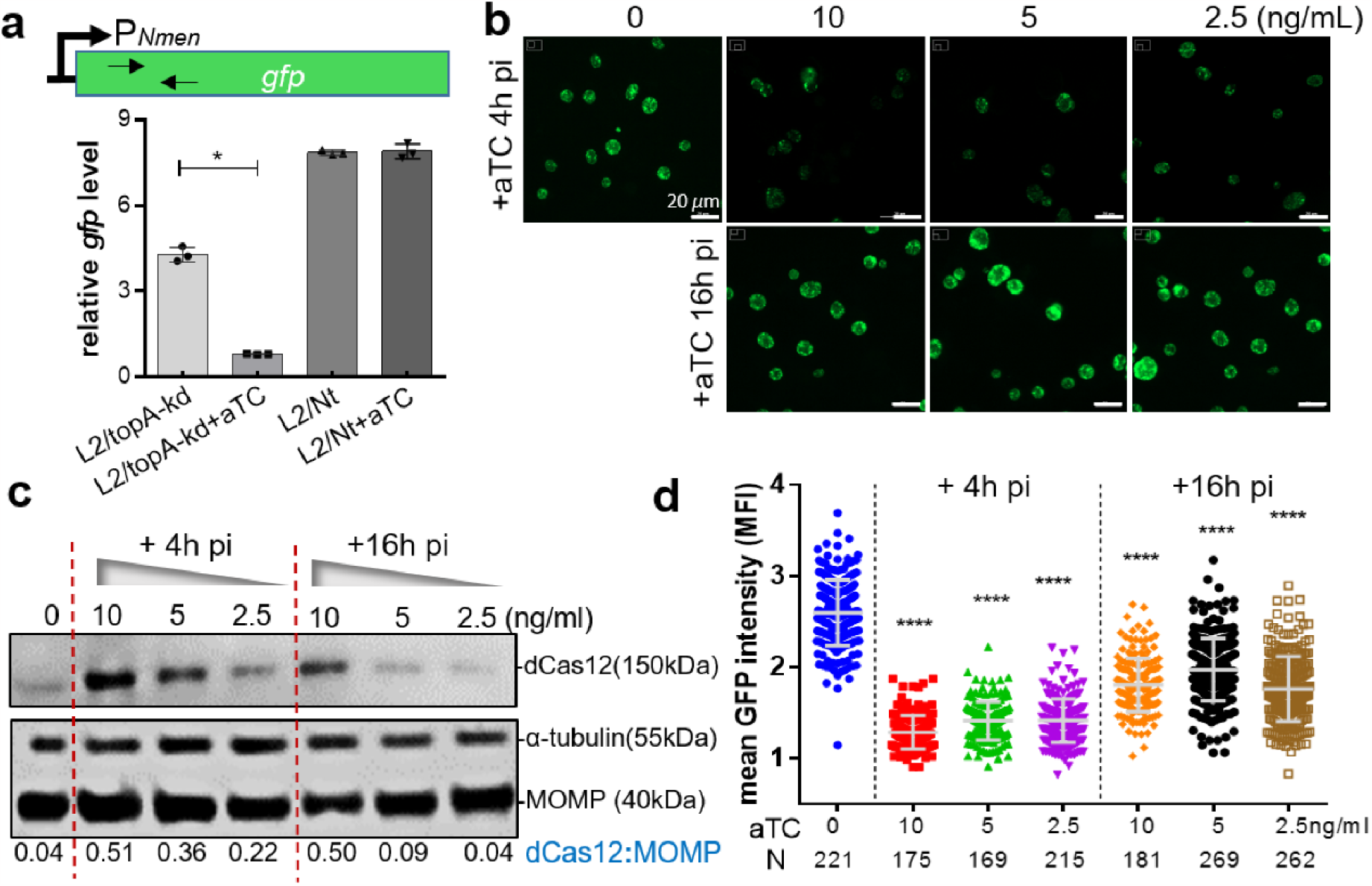
Dose- and time-dependent effects of targeted *topA* knockdown on P_*Nmen*_-*gfp* expression in *C. trachomatis* L2/topA-kd. (a) Quantification of *gfp* expression using RT-qPCR. The sites of primers used to detect *gfp* from the sample cDNA are indicated. The *gfp* mRNA concentrations were normalized to the DNA control as determined by qPCR targeting *tufA* and presented as mean ± SD of three biological replicates. (**b**) Live-cell images of *C. trachomatis*. HeLa cells were infected with *C. trachomatis* L2/topA-kd at MOI∼0.3 and cultured in aTC free medium. Increasing concentrations of aTC (0, 2.5, 5, or 10ng/mL) were added starting at 4 h pi or 16 h pi. The automated imaging acquisition was performed at 24 h pi under the same exposure conditions with Cytation 1. Scale bar=20 μm. (**c**) Immunoblotting analysis of dCas12 and MOMP expression. Increasing concentration of aTC was added at 4 h pi to induce dCas12 expression. Densitometry of the blot was assessed using ImageJ. Values are presented as the density of the dCas12 band (the upper panel) normalized to the MOMP band (the lower panel) from the same sample. Host cell α-tubulin was used as protein loading control. Data were collected from two independent experiments. Note: a small amount of dCas12 leaky expression was detected in the absence of aTC. (**d**) Measurement of GFP MFI (mean fluorescence intensity) in *C. trachomatis* infected cells grown in the absence or presence of aTC. Individual inclusions were analyzed using the Gen5 software. The MFI values are presented as mean ± SD from the indicated inclusion numbers (N) per condition in replicate wells. \*\*\*\**p* < 0.0001, comparison was made using one-way ANOVA followed by Tukey’s post-hoc test.

We next determined to what degree repression of *topA*, measured as a function of dCas12 expression, altered P_*Nmen*_-*gfp* expression in *C. trachomatis*. This is worthwhile because robust P_*Nmen*_-*gfp* expression could serve as a tool to visually and quantitatively monitor chlamydial growth (28, 29). *C. trachomatis* L2/topA-kd infected cells grown in the presence of increasing concentrations of aTC were used to assess the mean fluorescence intensity (MFI) of GFP using quantitative microscopy. This assay is based on automated live cell imaging in combination with green fluorescence and bright light detection. The intensity ratio of GFP to bright field in an individual inclusion was calculated to estimate the GFP MFI. Immunoblotting analysis of respective cell lysates was performed to quantify the levels of dCas12 and the major outer membrane protein (MOMP) for normalization.

When aTC (at the concentrations from 0 to 10 ng/mL) was added to L2/topA-kd cultures starting from 4 h pi, we observed the dose-dependent effects of *topA* repression: the larger aTC dose used, the more dCas12 produced, and the smaller GFP MFI detected (Fig. 3b-d). The decreased GFP in L2/topA-kd was consistent with the reduced EB yields (Fig. 2). Addition of the same amounts of aTC at 16 h pi had less effects on dCas12 induction along with slightly weakened GFP levels. In contrast, no significant changes in GFP levels were observed in strain L2/Nt (suppl. Fig. S1). These results indicate that the aTC dose, as well as the time of addition and its duration used, affects the expression level of dCas12 that is directly linked to the degree of *topA* repression in L2/topA-kd. Because RBs begin to asynchronously differentiate to EBs at ∼16 h pi and extant RB and EB forms co-exist at later time points, the differences in the time-related efficiency of dCas12 induction suggest that the growth rate or physiological state of *C. trachomatis* plays a role in determination of dCas12 induction. For example, it is unlikely that EBs express dCas12 to any great degree given their reduced metabolic activity and macromolecule synthesis (30, 31). Alternatively, the chlamydial developmental forms may have different responses to CRISPRi-induced *topA* repression. Thus, in addition to the effects related to EB yield and inclusion expansion, changes in P_*Nmen*_-GFP levels allow for sensitive detection of *topA* repression-mediated inhibition of *C. trachomatis* growth.

### Targeted *topA* knockdown interrupts RB-to-EB differentiation

It is important to determine what process of chlamydial development is impaired by *topA* repression. Because the IFU assay is unable to assess noninfectious forms, such as RBs, we performed real-time qPCR to quantify genomic DNA (gDNA) of *C. trachomatis* with primers specific to the coding region of *tufA*, encoding translation elongation factor EF-Tu (32). With similar infection rates in HeLa cells, the *C. trachomatis* strains, L2/topA-kd and L2/Nt, exhibited a similar increase in gDNA at 15 h pi under dCas12-inducing condition, suggesting significant levels of RB multiplication. There was no change in gDNA amounts at 24 h pi (Fig 4a).

**Figure 4.**
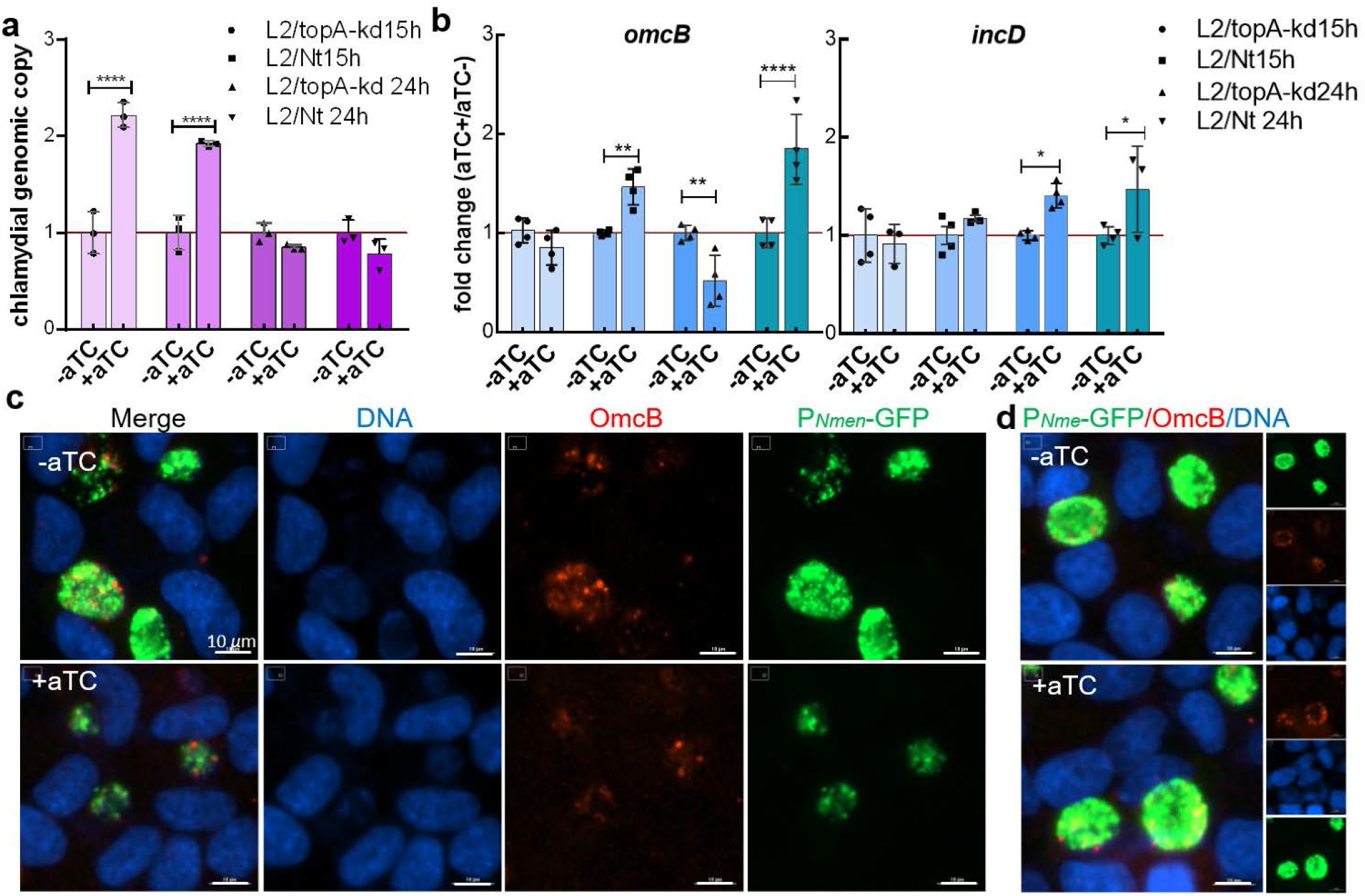
Secondary differentiation of RB to EB is impaired by *topA* knockdown in L2/topA-kd. (**a**) Analysis of *C. trachomatis* genomic copy numbers (i.e., gDNA) in the absence or presence of aTC using real-time qPCR targeting the *tufA* gene. Values are presented as the ratio of chlamydial DNA copy numbers per ng DNA in +aTC sample to that of -aTC sample, which is set at 1. Triplicate results in a representative experiment are shown. At least two independent experiments were performed. (**b**) Quantification of transcripts of *omcB* or *incD* in *C. trachomatis* using RT-qPCR. The mRNA concentrations were normalized to the DNA control as determined by qPCR targeting *tufA* and presented as mean ± SD of four biological replicates. (**c**)-(**d**) Immunofluorescent micrographs of *C. trachomatis* expressing OmcB. HeLa 229 cells were infected with *C. trachomatis* L2/topA-kd (**c**) or L2/Nt (**d**), cultured in the absence (-aTC,) and presence of aTC (+aTC, 10 ng/mL) for 20 h pi, and fixed at 24 h pi for IFA. Cells were immunolabeled with rabbit polyclonal antibody to *C. trachomatis* OmcB and visualized with Alexa Fluor 568-conjugated goat anti-rabbit antibody. DAPI*-*counterstained DNAs (blue) and *C. trachomatis* organisms expressing GFP (green) and OmcB (red) were shown. Scale bar= 10 μm. Statistical significance in all panels was determined by one-way ANOVA followed by Tukey’s post-hoc test. \**p* < 0.05, \*\**p* < 0.01, \*\*\*\**p* < 0.0001.

The lack of a meaningful difference in chlamydial gDNA amounts between L2/topA-kd and L2/Nt suggests that both strains progress through early developmental stages without issue. During infection, temporal expression of genes is correlated to the chlamydial developmental cycle (26, 33, 34). If perturbation of EB formation, but not RB replication, occurs, then *de novo* synthesis of early gene products is unaffected whereas late gene expression is diminished. We next tested whether we could find evidence of impaired EB formation (i.e. secondary differentiation). Nucleic acid samples were collected from HeLa cells infected with the L2/topA-kd or L2/Nt at 15 and 24 h pi. Expression of four genetic markers specific to *Chlamydia* development was examined using RT-qPCR: (i) *incD* encoding inclusion membrane protein, IncD, which is an early protein needed to establish the inclusion niche (35), (ii) *euo* encoding EUO that can bind to and repress late promoters (36, 37), (iii) *omcB* encoding a late 60kDa cysteine rich outer membrane protein OmcB, and (iv) *hctB* encoding histone-like protein 2, HctB (38). Regardless of whether aTC was added, less than ∼1.5 fold changes in transcript levels of *incD* and *euo* in L2/topA-kd and L2/Nt were detected at both 15 and 24 h pi (Fig. 4 and Fig. S2). In contrast, the addition of aTC decreased the transcript levels of *omcB* and *hctB* by ∼50% in L2/topA-kd at 24 h pi, while there was abundant transcription of *omcB* and *hctB* in L2/Nt. The disparity in OmcB expression between L2/topA-kd and L2/Nt was further confirmed by IFA (Fig. 4c). OmcB serves as a reliable EB marker because it provides integrity to the outer envelope via disulfide crosslinks with OmcA and MOMP (39). The signal of OmcB is much weaker in L2/topA-kd upon *topA* repression than mock repression. No decrease in immunolabeled OmcB was found in L2/Nt (Fig. 4d). The downregulated *hctB, omcB*, and OmcB expression, as well as continued *incD* and *euo* transcription together with the growth curve data, imply that *topA* knockdown disrupts the secondary differentiation from RB to EB. Conversely, the genomic DNA results suggest that RB replication is unaffected.

### Complementation of *topA* corrects *C. trachomatis* growth defect during CRISPRi-mediated knockdown

To confirm that the impaired growth phenotype observed is due to *topA* knockdown, we created a strain to complement TopA expression during knockdown as demonstrated previously for other targets (20, 21, 40, 41). The full-length *topA* gene with a six-histidine (His_6_)-tag was cloned and transcriptionally fused 3’ to the *dCas12* in pBOMBL12CRia(topA)::L2, resulting in pBOMBL12CRia-topA_6xH(topA)::L2 (Fig. 5a). In this vector, the *topA-His*_*6*_ is also under the control of the aTC-induced P_*tet*_ promoter. Thus, the *topA-His*_*6*_, as a source of the *topA*, is co-expressed with the *dCas12* that can be induced by addition of aTC. Because the *topA*-specific crRNA targets the promoter region of *topA* on the chromosome and causes growth defects, the functional product of *topA-His*_*6*_ should restore the normal growth of *C. trachomatis*.

**Figure 5.**
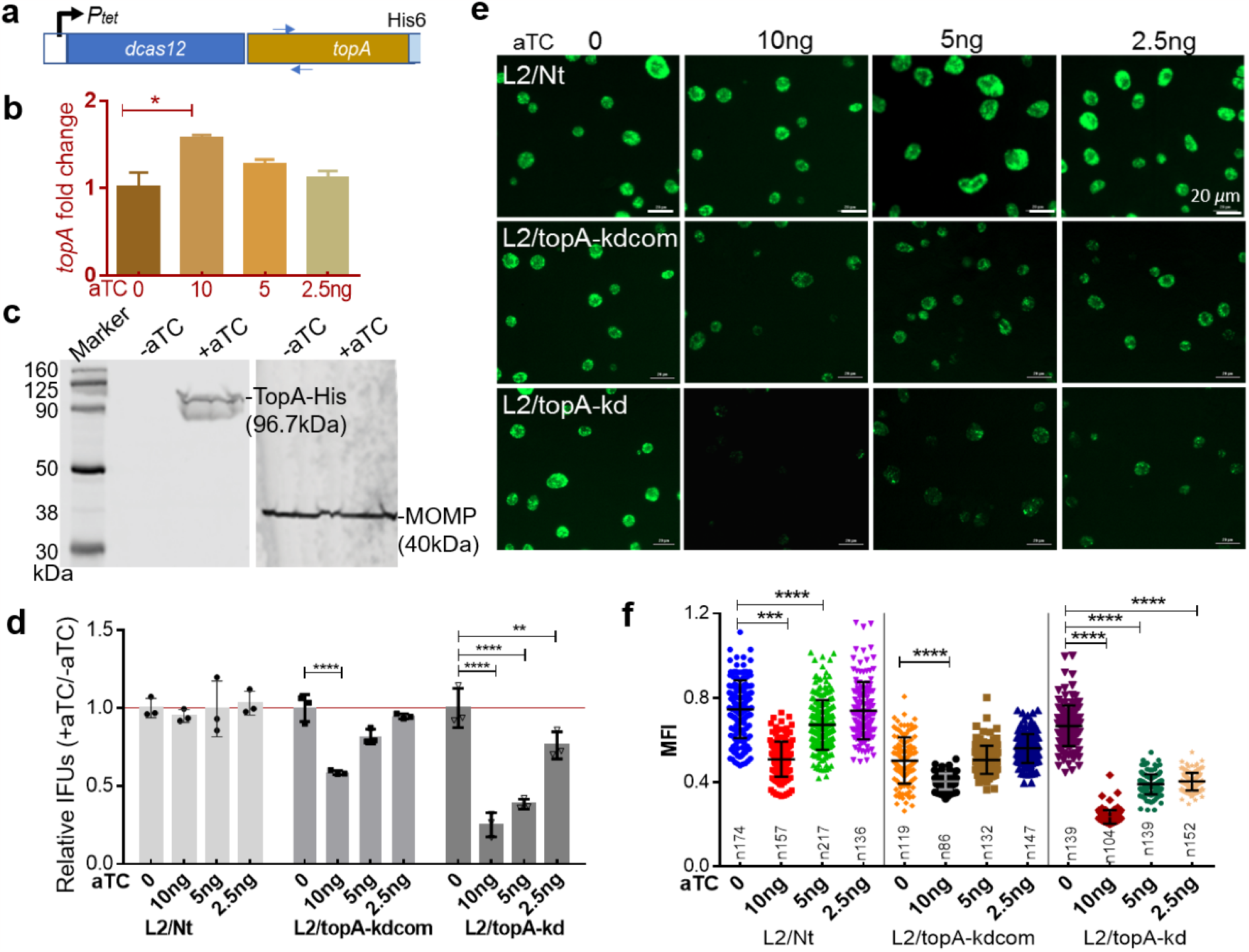
Complementation of the growth defect of *topA* knockdown in *C. trachomatis* by co-expressing *topA-His*_*6*_. (**a**) Schematic map of the expression vector containing *topA-His*_*6*_ that is co-regulated with the *dcas12* by P*tet*. (**b**) RT-qPCR analysis of *topA* transcripts in *C. trachomatis* L2/topA-kdcom. Nucleic acid samples from HeLa cells infected with L2/topA-kdcom were collected at 24 h pi. The locations of primers used to detect *topA* from the cDNA samples are shown in (**a**). (**c**) Immunoblotting displays the inducible expression of TopA-His_6_ in *C. trachomatis*. TopA-His_6_ protein from the lysates of *C. trachomatis* infected cells were isolated by 10% sodium dodecyl sulfate–polyacrylamide gel electrophoresis for immunoblotting with antibody against His_6_ or MOMP as a protein loading control. (**d**) Enumeration of EBs. *C. trachomatis* L2/Nt, L2/topA-kdcom, or L2/topA-kd infected cells were cultured for 40 hrs in the presence of increasing aTC amounts (at 0, 2.5, 5, and 10 ng/mL) and used for IFU assays. Values are presented from triplicate results in a representative experiment and are shown as mean ± SD. At least four independent experiments were performed. (**e**) Live-cell images of *Chlamydia* infected HeLa cells. Images were taken at 24 h pi with a 20× objective using Cytation 1. Scale bar=20 μm. (**f**) Analysis of changes in P_*Nmen*_-GFP levels as indicated as mean fluorescence intensity (MFI). Individual chlamydial inclusions from (**e**) were measured and calculated using Gen 5 software. The inclusion numbers (n) measured per condition are as indicated. For all panels, comparison was performed by ANOVA. \*\*\*\**p*<0.0001; \*\*\**p*<0.001, \*\*\*\**p* < 0.0001.

To determine whether plasmid-encoded *topA-His*_*6*_ could be conditionally expressed, *C. trachomatis* was transformed with pBOMBL12CRia-topA_6xH(topA)::L2. The resultant strain, L2/topA-kdcom, was used to infect HeLa cells. We confirmed the inducible expression of *topA* using real time RT-qPCR (Fig. 5b). As we did not have access to specific antisera against *C. trachomatis* TopA, antibody against His_6_ was used to examine the TopA-His_6_ protein by immunoblotting analyses of cell extracts in L2/topA-kdcom. Fig. 5c shows that an immunoreactive ∼97 kDa band was found in the presence of aTC. An additional ∼85 kDa immunoreactive band was seen. The nature of this band is uncertain, but we speculate that it could represent translation of an aborted *topA* transcript or an N-terminal degradation product of TopA-His_6_. The dCas12 expression was also detected by IFA (Fig. S3), indicating induction of TopA-His_6_ did not impair dCas12 expression in L2/topA-kdcom.

To adequately establish the optimal conditions for *topA* complementation, the growth of *C. trachomatis* was assessed by measuring the EB yields and GFP levels of the inclusions in the presence of increasing amounts of aTC (Fig. 5d-f). Interestingly, *C. trachomatis* L2/topA-kdcom displayed a different growth phenotype from that of L2/topA-kd or L2/Nt in response to aTC addition. In the presence of aTC, L2/topA-kd suffered from poor growth due to *topA* repression and L2/Nt did not, as expected. Under the same condition, L2/topA-kdcom accumulated EBs and had stronger GFP signal than L2/topA-kd, indicating its improved growth. However, the larger amount of aTC added, the smaller improvement was seen in L2/topA-kdcom. A possible explanation for these results is that excessive TopA could be toxic, so TopA-His_6_’s real action in correction of the growth defect is masked. Because L2/topA-kdcom carries a pBOMB4-based plasmid that is maintained at ∼5-7 copies per chromosome equivalent (42), a bacterial cell is expected to produce more than one copy of *topA*. The detrimental effect of an increased amount of TopA on bacterial growth has previously been reported (43). Nevertheless, the appropriate complementation of *topA* could restore EB yield to wild-type levels when aTC was used at ≤5 ng/mL, which was also the condition that hindered the growth of L2/topA-kd and had no inhibitory effects on the growth of L2/Nt.

Genetic complementation studies with plasmid-encoded *topA*-*His*_*6*_ demonstrate the need for appropriate inducible expression of TopA-His_6_ to restore the growth phenotypes to the levels compatible to WT *C. trachomatis*. These data fulfill molecular Koch’s postulates and further validate that CRISPRi-induced *topA* knockdown is responsible for the growth defect in the *C. trachomatis* L2/topA-kd strain.

### Targeted *topA* knockdown has pleiotropic effects on expression of DNA gyrase genes

Our data thus far reveal that CRISPRi-mediated *topA* repression has a profound impact on *C. trachomatis* growth along with downregulated transcription of chromosomal genes (i.e., *omcB* and *hctB*) and the plasmid-encoded P_*Nmen*_-*gfp*. These data are in line with previous studies in *E. coli* that suggested a role of TopoI in controlling transcription processes (2, 44-46). We sought to address whether *topA* repression altered transcription of gyrase encoded genes (*gyrA/gyrB)* and TopoIV encoded genes (*parE/parC)*, given a potential feedback mechanism of Topo gene regulation reported by Orillard and Tan (19). In the *C. trachomatis* chromosome, *ctl0443/ct191* encoding a hypothetical protein is adjacent to *gyrB* and *gyrA*, and there is a putative promoter upstream of *ctl0443/ct191* (Fig. 6). Located at a different locus, *parE* and *parC* are also adjacent genes. These polycistronic mRNAs were quantified using real-time RT-qPCR with *C. trachomatis* infected HeLa cells harvested at 24 h pi.

**Figure 6.**
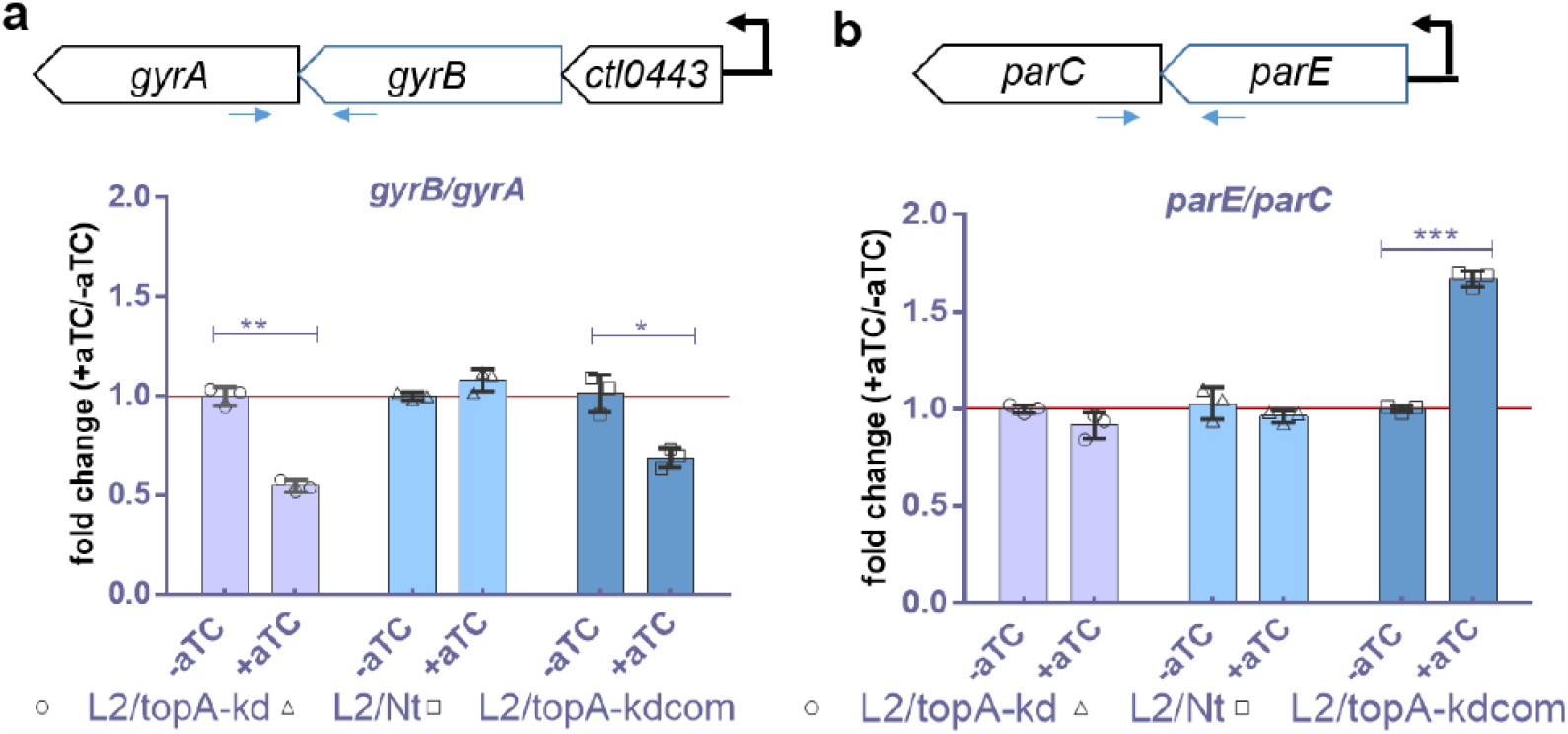
The effects of CRISPRi-induced *topA* knockdown on expression of DNA gyrase genes and topoIV genes in *C. trachomatis*. (**a**) Schematic map of *gyrB/gyrA* operons in *C. trachomatis* and detection of their transcript products using RT-qPCR. (**b**) Schematic map of *parE/parC* in *C. trachomatis* and detection of their transcript products using RT-qPCR. L2/top-kd infected HeLa cells grown in the presence or absence of aTC were harvested at 24 h pi for total RNA preparation and then cDNA synthesis. Results of a representative experiment from triplicate samples are reported as mean ± SD. Three independent experiments were performed. \*\**p* < 0.005, *** P< 0.001. Comparison was made using one-way ANOVA and Tukey’s post-hoc test. Primer pairs overlapping gene pairs used for RT-qPCR analysis are shown (arrows).

Steady-state levels of *gyrB/gyrA* transcripts were unchanged in L2/Nt in the presence of aTC, while the *gyrB/gyrA* transcripts were reduced in L2/top-kd and L2/topA-kdcom (Fig. 6). These results are unsurprising as the different bacterial Topos possibly compensate for a defect in one enzyme by varying expression of another as observed in *E. coli* (8). Unlike the results for *gyrB/gyrA*, expression of *parE/parC* was unchanged in L2/top-kd and L2/Nt and was increased (∼1.7 fold) in L2/topA-kdcom. Since addition of aTC induced CRISPRi-mediated *topA* repression in L2/topA-kd, these results imply that that the proportion of gyrase synthesis was regulated in *C. trachomatis* in response to *topA* repression under our testing conditions. Reduced *gyrB/gyrA* expression in L2/topA-kd is not due to mutation in the promoter region upstream of *ctl0443/ct191* as analyzed by PCR and DNA sequencing analysis (data not shown). For L2/topA-kdcom, in which *topA-his6* was induced by adding aTC, increased TopoIV and decreased gyrase transcripts were measured. These data provide partial explanations for the detected differences in the growth and molecular phenotypes between L2/Nt, L2/topA-kd, and L2/topA-kdcom (Figs. 2-5). The observations showing that changes in *topA* levels in *C. trachomatis* can trigger alterations in type II Topo expression are consistent with the view that Topo activity in bacteria is carefully balanced to sustain DNA supercoiling levels.

### Targeted *topA* knockdown affects the response of *C. trachomatis* to moxifloxacin

A low level of tolerance to quinolones was associated with reduced expression of quinolone targets, gyrase and/or TopoIV, in bacteria (8, 47). Having found decreased transcription of *gyrB/gyrA* in L2/topA-kd due to *topA* repression, we postulated that the sensitivity of *C. trachomatis* to moxifloxacin (Mox) may be altered. To test this hypothesis, the minimal inhibitory concentration (MIC) of Mox was initially determined with *C. trachomatis* reference strain, L2/434/Bu, in HeLa cells. The concentrations of Mox that resulted in decreases in IFUs to 50% and 99% of the unexposed culture were 4.5 ng and 50 ng per mL, respectively, using IFU assay and IFA (Fig. S4). The MIC was determined as 100 ng/mL, where no IFUs were detected in subculture. Similar results were obtained in *C. trachomatis* strain L2/Nt (with wild-type *topA*).

We next sought to determine if *topA* knockdown influenced the sensitivity of *C. trachomatis* to Mox at a sub-MIC (5 ng/mL =1/20 MIC). Sub-MIC was used because we were interested in changes in the growth phenotype in live *C. trachomatis*. The growth patterns of *C. trachomatis* L2/topA-kd, L2/Nt, and L2/topA-kdcom in HeLa cells were assessed in the absence of Mox, presence of Mox alone, or aTC+Mox by measuring the inclusion size and IFUs.

The addition of Mox or aTC+Mox reduced inclusion sizes in all strains tested (Fig. 7a-b). Interestingly, Mox or aTC+Mox decreased GFP levels in L2/Nt and L2/topA-kdcom, while aTC+Mox, but not Mox alone, weakened the GFP signal in L2/topA-kd (Fig. 7c). Consistent with the reduced inclusion sizes induced by Mox, EBs in L2/topA-kd and L2/Nt were ∼30% and ∼50% less than the untreated controls, respectively, at 40 h pi. Conversely, there was no significant change in L2/topA-kdcom. However, with aTC+Mox in the medium, EB yields were decreased in all strains with the most prominent reduction in L2/topA-kd (∼90% less) as analyzed by IFU assay. In support of the IFU data, decreased chlamydial gDNA yields were measured at 24 h pi for L2/topA-kd and L2/Nt in the aTC+Mox condition. Interestingly, the gDNA content in L2/topA-kdcom was unchanged by either Mox or aTC+Mox. We confirmed the induction of dCas12 expression in L2/topA-kdcom by addition of aTC+Mox (Fig. S5), suggesting concurrent *topA* expression, which was co-transcribed with dCas12, and the CRISPRi-mediated *topA* repression under these conditions (see also Fig. 5). The unchanged gDNA content, in contrast to the reduced EBs in L2/topA-kdcom, imply the continued replication of RBs without progression through the developmental cycle, the induction of persistent forms that are viable but non-infectious, or, potentially, dead bacteria.

**Figure 7.**
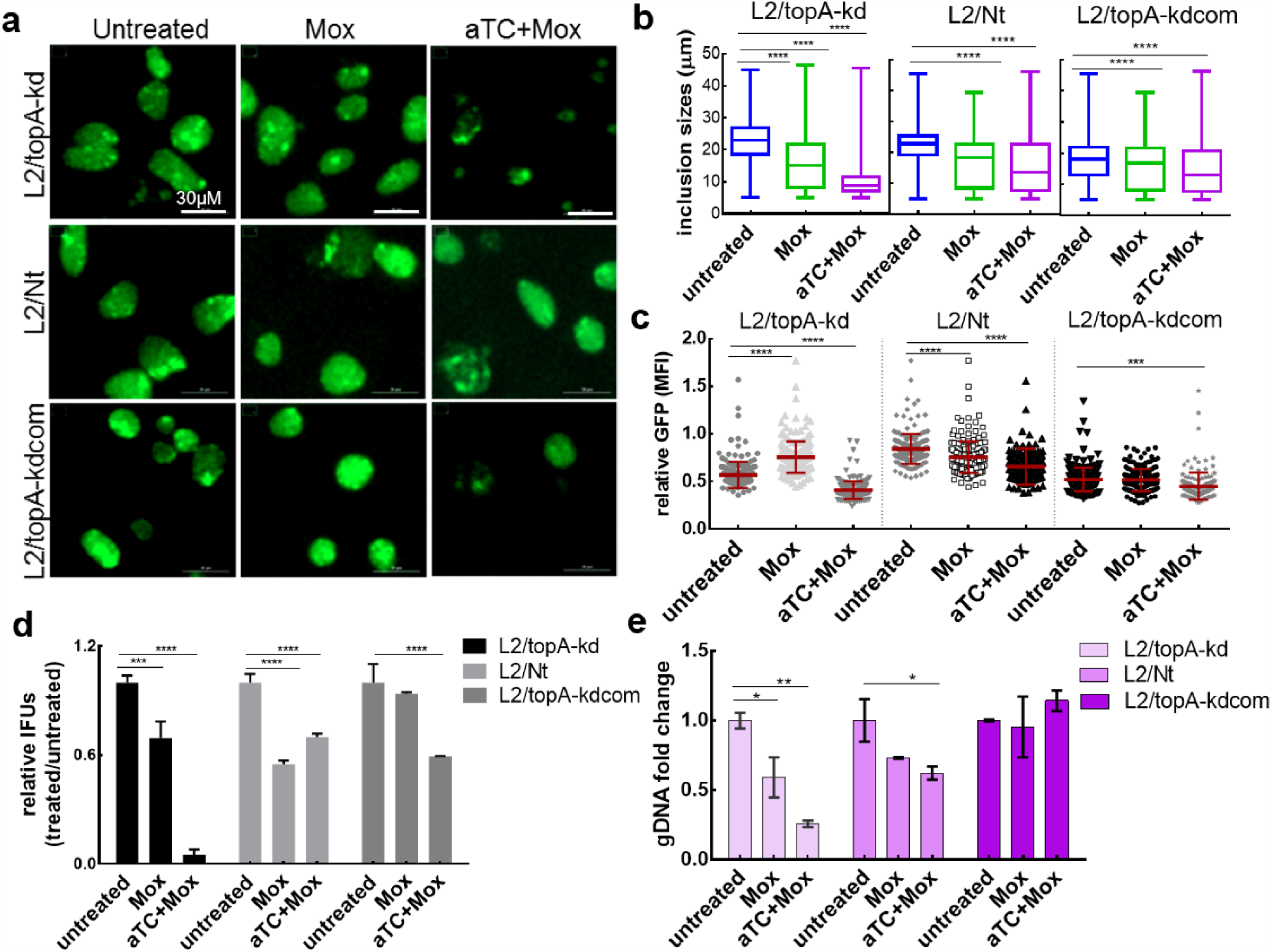
Analysis of the response of *C. trachomatis* to the antibiotic moxifloxacin. (**a**) Live-cell images of *Chlamydia* infected HeLa cells. *C. trachomatis* L2/topA-kd, L2/Nt, or L2/topA-kdcom infected cells were cultured in the absence or presence of Mox or aTC+Mox and imaged at 44 h pi. Scale bar=30μm. (**b**)-(**c**) Comparison of the chlamydial inclusion sizes (**b**) and the GFP MFI (**c**) of L2/topA-kd to those of L2/Nt and L2/topA-kdcom. Two hundred individual chlamydial inclusions per condition from images in (**a**) were measured using Gen 5 software. **d**. Enumeration of EB yields using IFU assay. *C. trachomatis* infected cells were harvested at 40 h pi for IFU assay. Values are presented from triplicate results in a representative experiment and are shown as mean ± SD. Three independent experiments were performed. (**e**) Analysis of *C. trachomatis* gDNA in the presence or absence of aTC using real-time qPCR. Values are presented as the ratio of chlamydial DNA copy numbers per ng DNA in treated sample to that in the untreated sample, which is set at 1. Triplicate results in a representative experiment are shown. Three independent experiments were performed. Statistical significance in all panels was determined by one-way ANOVA followed by Tukey’s post-hoc test. \**p* < 0.05, \*\**p* < 0.01, \*\*\*\**p* < 0.0001.

Together, these results imply disparities in the responses of *C. trachomatis* to the sub-MIC of Mox between L2/topA-kd, L2/Nt, and L2/topA-kd, suggesting that the levels of *topA* expression influences the chlamydial response to moxifloxacin.

## Discussion

Topos have been extensively studied since the first discovery of bacterial Topo I in 1971 (48). However, the understanding of Topos and their relevance to *C. trachomatis* development has only slowly progressed, hampered in large part by the lack, until recently, of tractable genetic techniques for *Chlamydia*. The recent development of CRISPRi as a genetic tool to inducibly repress transcription in *Chlamydia* allowed us to demonstrate an indispensable role of TopA in controlling *C. trachomatis* developmental progression. Our studies have characterized the growth phenotype and selected developmentally expressed genes in *C. trachomatis* following *topA* repression. We also evaluated the consequences of *topA* repression for the bacterial cells in response to moxifloxacin (targeting gyrase) and showed that *Chlamydia* displays enhanced sensitivity to moxifloxacin when *topA* is repressed. These results strongly support the notion that TopA functions together with the gyrase to support the developmental process of *C. trachomatis* as illustrated in Figure 8. The ability to conditionally manipulate essential genes through CRISPRi will allow an assessment of fundamental transcriptional and replication states of the *C. trachomatis* genome.

**Figure 8.**
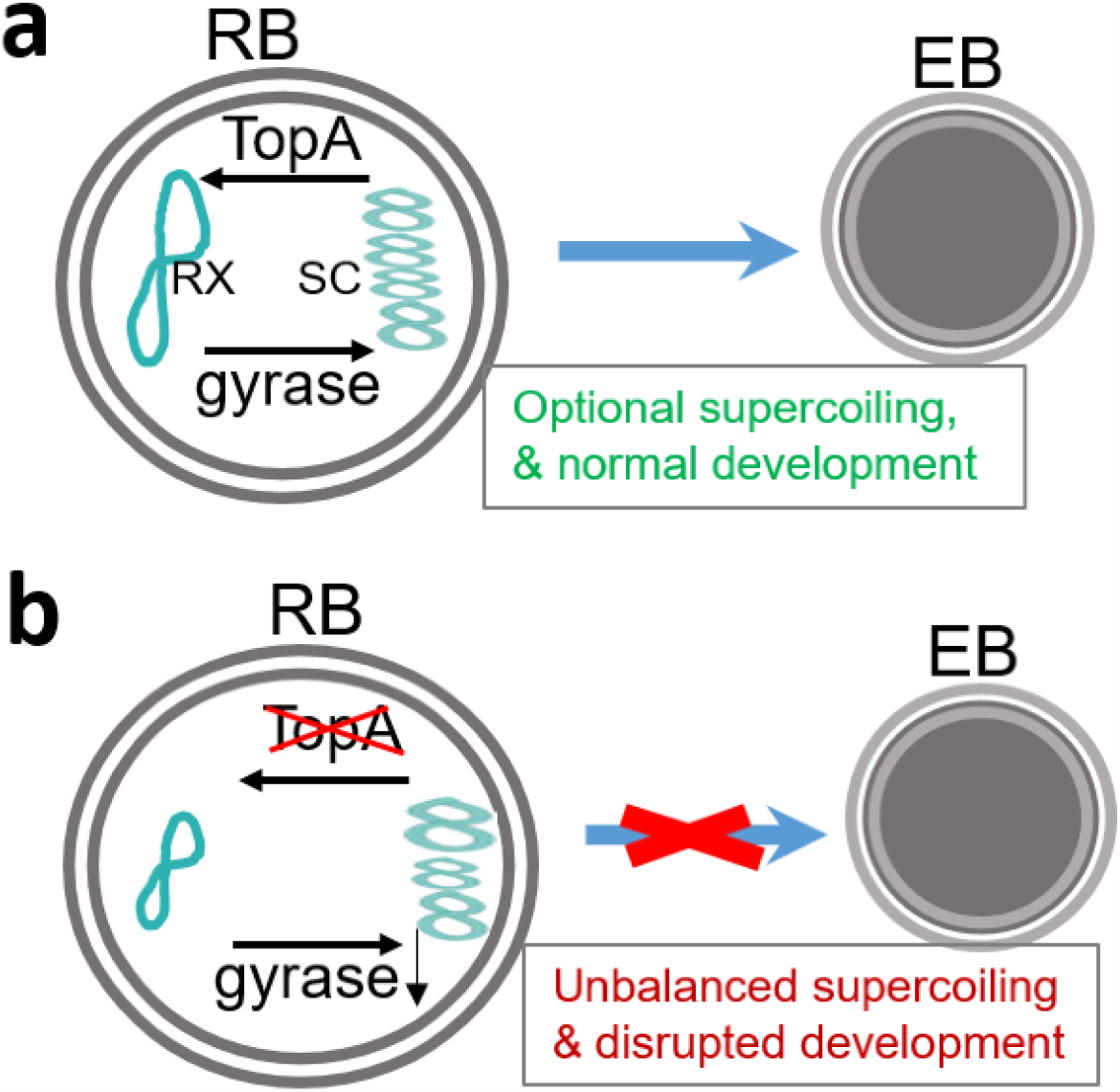
Schematic highlighting the role of TopA in *C. trachomatis* developmental cycle. **a**. In wild-type *C. trachomatis*, optimal supercoiling levels during chlamydial developmental cycle progression is maintained by action of both *topA* that relaxes DNA supercoiling (RX) and gyrase that induces negative supercoiling (SC). **b**. When *topA* is repressed, the DNA supercoiling is predicted to increase, resulting in changes in expression of supercoiling-sensitive genes (e.g., chromosomal *gyrB/gyrA* and *omcB*, and the plasmid-encoded P_*Nmen*_-*gfp*), thus, perturbing chlamydial development. Our data indicate that the carefully balanced activities of TopA and gyrase contribute to the completion of the chlamydial developmental cycle.

### Utility of CRISPRi system in *C. trachomatis*

Studying essential genes, such as Topo encoded genes, is challenging because conventional gene disruption strategies lead to lethality. New technologies such as CRISPRi can circumvent such obstacles. CRISPRi has provided inducible knockdown of gene expression and enabled genetic approaches to studying essential gene function in several bacterial pathogens (49-51). Only a handful of studies have reported the use of CRISPRi in intracellular bacteria (20, 52, 53). We have demonstrated successful knockdown of *topA* in *C. trachomatis* using a plasmid-based CRISPRi system that relies on the combination of inducible dCas12 and a *topA*-specific crRNA. The incorporation of P_*Nmen*_-GFP as a reporter greatly facilitates quantifying and monitoring the influence of *topA* repression on *C. trachomatis* growth. Moreover, we show the effects of dose, time, and duration of aTC addition on the efficiency and degree of CRISPRi-mediated *topA* repression. These results reflect the *C. trachomatis* developmental cycle and the unique challenges of working with this pathogen. Since RBs are highly active in macromolecular synthesis and have a metabolism differing from EBs (31), it is unsurprising that the growth rate and physiological state of *C. trachomatis* play a role in determination of dCas12 induction as observed in the current studies. It is also likely that the distinct chlamydial forms respond to aTC and induced dCas12 differently, a factor that needs to be taken into consideration when interpreting the results.

A very important issue when using CRISPRi for gene knockdown is its specificity. Three independent observations indicate targeted *topA* repression is specifically mediated by CRISPRi in our experiments. First, according to our genome-wide sequence analysis, no sequence similarity to the *topA*-targeting crRNA was found in *C. trachomatis*. Second, the negative effects of *topA* repression on *C. trachomatis* were rescued by complementation with a plasmid-encoded *topA* when expressed at the appropriate level and induction time. Third, no reduction of *topA* transcription was observed when using a non-targeting crRNA that has no homology to chlamydial sequences.

### The role for TopA in bacteria-host interaction

Since TopA is an essential enzyme that primarily acts to relax negative supercoiling during transcription, we evaluated the effects of repression of this enzyme on *C. trachomatis* growth during infection in host epithelial cells. We demonstrated that *topA* knockdown profoundly influences the activities of *C. trachomatis* (Figs. 2-4). The data, with combined methods for detecting differences in growth between strains either having WT or repressed *topA*, indicate the requirement of TopA to complete the chlamydial developmental cycle. Surprisingly, however, we found that chlamydial growth of the L2/topA-kdcom strain was slowed when plasmid-encoded *topA*-*His*_6_ was highly expressed. We evaluated the possibility that the detrimental effect of an increased amount of TopA might be associated with the suboptimal growth. It was found that under the optimal conditions for *topA-His*_*6*_ induction, L2/topA-kdcom displayed improved growth (Fig. 6). Thus, the excessive plasmid-expressed TopA is likely unfavorable for *C. trachomatis* and only a suitable level of TopA-His_6_ can complement *topA* repression-related growth defects in *C. trachomatis*. The differences in *topA* levels between L2/topA-kd (*topA* repression) and L2/topA-kdcom (*topA* overexpression) is sufficient to explain their varied growth phenotypes in human epithelial cells.

### The role for TopA in bacterial gene regulation

DNA topology affects regulation of gene expression both globally and locally in bacteria. Recent genome-wide transcriptomic data suggest that elaborate mechanisms are employed by bacteria to coordinate transcription rates and Topo activity to adjust supercoiling levels in the promoter regions of differentially expressed genes (44, 46, 54). Previous studies suggested changes in DNA topology occur during the developmental cycle (11, 15, 16, 18) and that several putative promoters and respective genes appeared to be more “supercoiling-sensitive” than others in *C. trachomatis* (16, 18, 19). For example, all three promoters of chlamydial Topo genes act in a supercoiling-dependent manner. Complementing and extending these findings, our data demonstrate that the transcript levels of gyrase genes were decreased following *topA* repression in *C. trachomatis*, while the levels of TopoIV were unchanged. We did not have access to specific antibodies against the subunits of chlamydial gyrase and TopoIV and thus did not measure their levels directly; however, decreases in the levels of *gyrA/gyrB* transcripts suggest that the levels of the gyrase holoenzyme were likely reduced. In *E. coli*, TopoI relaxes negatively supercoiled DNA and has been shown to sustain the steady-state level of supercoiling by balancing the activity of DNA gyrase (3, 5). Since the main functions of Topos are to prevent excessive supercoiling that is deleterious for bacterial cells, we speculate that, if *topA* is knocked down, the enzyme will no longer relax supercoiling. Subsequent decreases in gyrase expression likely occur, perhaps, to balance the supercoiling levels for maintaining *Chlamydia* survival (Fig. 8). Although the current study did not directly demonstrate increased DNA supercoiling in *Chlamydia* after *topA* repression, increases in general DNA supercoiling were observed upon topoI inhibition in *E. coli* (5) and *S. pneumoniae* (55), and the opposite was observed by gyrase inhibition using novobiocin. Our data recapitulate the relationship between TopA and the type II Topos and the view that opposing catalytic activities of TopA (to relax) and DNA gyrase (to supercoil) ensure homeostasis of chromosomal and plasmid DNA supercoiling.

The correlation was also established between *topA* repression and the altered expression of chromosomal genes (i.e., *omcB*, and *hctB*) and the plasmid-encoded P_*Nmen*_-*gfp* in strain L2/topA-kd in comparison with the L2/Nt. According to previous studies, early genes (i.e. *incD* and *euo*) and the late gene *omcB* were supercoiling insensitive *in vitro*. We demonstrate that expression levels of *incD* and *euo* were unchanged, while *omcB* transcription was inhibited when *topA* was repressed. These results indicate that *omcB* may sense DNA topological levels in *C. trachomatis*. The differences in *omcB* levels in response to supercoiling between the current study and previous reports might reflect that many *C. trachomatis* promoters or genes are likely responsive to supercoiling in a context-dependent manner *in vivo*. The role of DNA supercoiling in regulating *hctB* transcription has not been defined yet as its supercoiling sensitivity remains to be determined. However, it was shown that *hctA* encoding the histone-like protein 1 was supercoiling insensitive *in vitro* (16). Future studies will use RNA sequencing to determine global transcriptome changes in response to *topA* repression.

### Response of *C. trachomatis* to moxifloxacin

To evaluate a direct outcome linked to *topA* repression, we examined whether *topA* repression influenced the response of *C. trachomatis* to moxifloxacin. Although TopA is not the target of quinolones, inhibiting or overexpressing *topA* changes the expression levels of gyrase and TopoIV genes (e.g., Fig. 6). Unlike novobiocin, that inactivates ATPase activity of the GyrB subunit of gyrase, quinolones dually inhibit gyrase GyrA and Topo IV activities, forming a poisonous Topo-quinolone-DNA complex that eventually breaks double-stranded DNA leading to bacterial death (6, 7). Aminocoumarin and quinolones are potent inducers of SOS-related stress responses (43). Thus, changes during antibiotic exposure may reflect effects of both supercoiling and that of the supercoiling-independent stress response, and these mechanistically different contributors are hard to distinguish. For this reason, we used a sub-MIC concentration of moxifloxacin to determine the impacts of *topA* repression on moxifloxacin sensitivity. With the *C. trachomatis* strain L2/topA-kd, we were able to link the enhanced sensitivity to Mox with the repression of *topA*. This result is unexpected, as, based on observations from other bacteria, decreased levels of the drug target (i.e., gyrase) may reduce formation of poisonous complexes and thus weaken drug action, in turn leading to relative quinolone tolerance. On the other hand, moxifloxacin had different effects on the strain L2/topA-kdcom depending on the induction of *topA* levels. While this strain displayed a low level of tolerance to moxifloxacin under uninducing conditions (-aTC), it was inhibited under *topA* knockdown conditions (+aTC). This complexity observed is probably associated with the relative scale of *topA-his*_6_ overexpression and *topA* repression in L2/topA-kdcom. To distinguish these two opposing observations, we will need to evaluate *topA* overexpression in a background strain lacking CRISPRi elements. We cannot exclude the possibility that there are a proportion of dead *Chlamydia* in the presence of moxifloxacin when *topA* was highly overproduced or repressed. However, most bacteria remained viable under our testing conditions because they readily resume normal growth after aTC+Mox containing medium was replaced with normal medium starting at 24h pi (Shen et al unpublished observation).

Our data indicate that TopA is required for *C. trachomatis* development. There are still open questions. For example, how do the integrated activities of TopA, gyrase, and the TopoIV specifically contribute to the metabolism of *C. trachomatis*? This is an important question because Topos are drug targets for the development of new antibacterial therapy (56, 57). Although resistance to quinolones is currently rare in clinical *C. trachomatis* isolates, there were reports showing the potential of acquiring quinolone resistance *via* mutations in the *gyrA* gene after prolonged exposure to sublethal Mox concentrations in culture (58). In addition, mutations in *ygeD* encoding an efflux protein was associated with quinolone resistance in clinical isolates (59). Another question is whether endogenous plasmid genes, similar to the exogenous P_*Nmen*_-*gfp*, sense changes in Topo-mediated supercoiling *in vivo*. Supercoiling data obtained from the small endogenous plasmid will provide a good estimation of the relationship between DNA supercoiling and gene transcription on the chromosome. They will also improve our understanding of the pathogenesis of infection because the chlamydial plasmid is a central virulence factor (60, 61): plasmid-encoded Pgp4 regulates expression of plasmid and chromosomal genes, including the secreted glycogen synthase, GlgA. Despite the evidence that *topA* levels affect transcription of the selected *C. trachomatis* stage-expressed genes, the overall influence of TopA activity on the *C. trachomatis* plasmid and chromosome requires additional investigation. Future studies will attempt to evaluate *in vivo* the changes in DNA topology when altering Topo activity or expression levels as well as the relative contributions of gyrases and TopoIV in DNA supercoiling during the chlamydial developmental cycle.

## Materials and Methods

### Reagents and antibodies

Antibiotics and dimethyl sulfoxide (DMSO) were purchased from MilliporeSigma (St. Louis, MO, USA). Moxifloxacin stock solution was dissolved in 100% DMSO at 10 mg/mL. In all experiments, the Moxifloxacin stock was diluted in the corresponding culture medium, and controls lacking moxifloxacin were performed using an equal percentage of DMSO. FastDigest restriction enzymes, alkaline phosphatase, and DNA Phusion polymerase were purchased from ThermoFisher (Waltham, MA). The following primary antibodies were used: (i) a mouse monoclonal antibody (L2I-45) specific to the LGV L2 MOMP (48), (ii) a rabbit polyclonal anti-OmcB (kind gift from Tom Hatch, University of Tennessee), (iii) a rabbit polyclonal AsCpf1/Cas12a antibody (catalog #19984, Cell Signaling Technology), (iv) a rabbit polyclonal ant-His6 antibody (catalog #213204, Abcam), and (iv) a mouse monoclonal antibody to tubulin (catalog #T5168, MilliporeSigma). The secondary antibodies used were Alexa Fluor 568-conjugated goat anti-mouse IgG (catalog #A11004) from Invitrogen (Carlsbad, CA, USA) and horseradish peroxidase (HRP)-conjugated goat anti-rabbit IgG (catalog #213204, Abcam) and HRP-conjugated anti-mouse IgG (catalog A0168, MilliporeSigma).

### Cell culture and *C. trachomatis* infection

Human cervix adenocarcinoma epithelial HeLa 229 cells (ATCC CCL-2.1) were cultured in RPMI 1640 medium (Gibco) containing 5% heat-inactivated fetal bovine serum (Sigma-Aldrich), gentamicin 20 μg/mL, and L-glutamine (2 mM) (RPMI 1640-10) at 37°C in an incubator with 5% CO_2_. Cells were confirmed to be Mycoplasma-negative by PCR as described previously (62). *C. trachomatis* strains used were listed in Table 1. The strains were authenticated by sequencing of whole PCR product of *ompA* and by staining with antibody to the LGV L2 MOMP. Spectinomycin (500 μg/mL) and cycloheximide (0.5 μg/mL) were added to propagate transformed *C. trachomatis* strains. Stocks of WT and transformed *C. trachomatis* were made every one year and aliquots of purified EBs were stored in -80° C until use. For infection and *C. trachomatis* analysis, cells grown in 96-well plates (cat. No 655090; Greiner) were inoculated with isolated EBs with a dose that results in ∼30-40 % of cells being infected, centrifuged with a Beckman Coulter model Allegra X-12R centrifuge at 1,600 x g for 45 min at 37°C, and cultured in RPMI 1640-10 without cycloheximide at 37°C for various times as indicated in Results. Fresh medium was added to the infected cells and incubated at 37 °C for various time periods as indicated in each experimental result. For comparison, different strains were infected side-by-side in the same culture plate with a setup of at least triplicate wells pre condition.

### Plasmids and transformation

Plasmids and primers used in this study are listed in Tables S1 and S2 in the supplemental material. The spectinomycin resistance encoding empty vector plasmid, pBOMBL12CRia(e.v.)::L2 (aka pBOMBL-As_ddCpf1vaa::L2) (20), was digested with BamHI and treated with alkaline phosphatase. Two nanograms of the *topA*-targeting or non-targeting crRNA gBlock (Suppl. Table S2) were mixed with 25ng of the BamHI-digested pBOMBL12CRia(e.v.)::L2 in a HiFi reaction (NEB) according to the manufacturer’s instructions. The reaction mix (2 μL) was then used to transform 25 μL of chemically competent 10-beta cells (C3019H; NEB), which were subsequently plated on Luria-Bertani (LB) agar plates containing 100 μg/mL spectinomycin. Individual colonies were screened for the presence of the correct plasmids after miniprep (Qiagen kit) extraction from overnight cultures using restriction enzyme digest and Sanger sequencing. For the complemented vector, *topA-His*_*6*_ was PCR amplified using DNA Phusion polymerase, the primer pair (topA/(dCas12vaa)/5’ and topA_6xH/(pL12CRia)/3’) (Table S2), and *C. trachomatis* serovar L2/434/Bu genomic DNA as template. The PCR product was confirmed for correct size by agarose gel electrophoresis and purified using a PCR purification kit (Qiagen). The vector pBOMBL12CRia(*topA*)::L2 was digested with Fast-digest SalI and treated with alkaline phosphatase as described above. The purified PCR product of *topA-His*_*6*_ (13ng) was mixed with 25 ng of the SalI-digested pBOMBL12CRia(*topA*)::L2 in a HiFi reaction. *E. coli* 10-beta transformants were obtained and plasmids verified as described above. Two micrograms of sequencing verified CRISPRi plasmids were used to transform *C. trachomatis* serovar L2 lacking its endogenous plasmid (-pL2) as described previously (20, 24) and using 500 μg/mL spectinomycin as selection. DNA was extracted from chlamydial transformants to verify plasmid sequence.

### Microscopy analysis

Automated live-cell images in 96-well culture plates were acquired using an imaging reader Cytation1 (BioTek Instrument). Gen5 software was used to process and analyze the inclusion morphology (e.g., inclusion size, numbers, and mean fluorescence intensity (MFI)). For indirect immunofluorescence assay (IFA), the *C. trachomatis* infected cells were fixed with 4% (w/v) paraformaldehyde dissolved in PBS (pH 7.4) for 15 min at room temperature, permeabilized with 0.1% (v/v) Triton X-100 for an additional 15 min, and blocked with 2% (w/v) bovine serum albumin (BSA) in PBS for 30 min. Then, cells were incubated with the indicated primary antibody overnight at 4°C, followed by incubation with Alexa Fluor 488/568-conjugated secondary antibody for 45 min at 37°C. 4^′^,6-diamidino-2-phenylindole dihydrochloride (DAPI) was used to label DNA. In some experiments, cell images were visualized and photographed using an inverted fluorescence microscope (Zeiss Axio Observer D1) and analyzed with the AxioVision software, version 4.8.

### *C. trachomatis* enumeration and end point one-step growth curve

To evaluate infectious EB progeny, inclusion forming unit (IFU) assays were performed in 96-well plates. Briefly, *C. trachomatis* infected cells in culture plates were frozen at -80 °C, thawed, scraped into the medium, serial-diluted, and then used to infect a fresh monolayer of HeLa cells. The infected cells were cultured in RPMI 1640-10 with 500 μg/mL spectinomycin and without cycloheximide at 37 °C for 40 hrs. Cells were fixed, processed, and then stained with antibody against LGV L2 MOMP. Images were taken using fluorescence microscopy and the inclusion numbers in triplicate wells were counted. The total EB numbers are presented as the number of IFUs per mL. In some experiments, the IFU value was normalized to the control and presented as percentage. For growth curves, the cultures were harvested at the time-points 0, 12, 24, 30, and 48 h pi, and followed the same procedure as described above to titrate IFUs.

### Antimicrobial susceptibility testing

Minimal inhibitory concentrations (MIC) of antibiotics were tested in 96-well plates as described (63, 64). Briefly, purified EBs (10,000/well) were used to infect HeLa cell monolayers, followed by centrifugation with a Beckman Coulter model Allegra x-12R centrifuge at 1,600 x g for 45 min at 37°C. After the removal of supernatant, the infected cells were washed with phosphate buffered saline (PBS) once and cultured in RPMI-10 containing the appropriate concentration of test antimicrobials in a volume of 100 μL at 37°C in a humidified incubator with 5% CO_2_ for various time periods as indicated in each experimental result. *C. trachomatis* inclusions were immunolabeled with anti-MOMP antibody and enumerated using fluorescence microscopy. The MIC was defined as the lowest concentration of drug without visible *C. trachomatis* growth in the subculture.

### Nucleic acid analysis

For nucleic acid preparation, *C. trachomatis* infected HeLa cells in 24-well plates were harvested at 15 and 24 h pi, respectively. Quick DNA/RNA miniprep kit (catalog # D7001, Zymo Research) was used to isolate DNAs and RNAs sequentially as instructed by the manufacturer. Residual DNA in the RNA samples was removed by treatment with 20U RNase-free DNase I in-column for 30 min at room temperature and extensively washing. A total of 2 μg of RNA per sample was reverse transcribed into cDNA using the high-capacity cDNA reverse transcriptase kit (Catalog # 4368814, Applied Biosystems). The Fast SYBR green master mix (Applied Biosystems) was used for qPCR assay in 20 μL of reaction mixture on a real-time PCR system (Bio-Rad) with the primer pairs listed in Table S2. Each sample was analyzed in triplicate in a 96-well plate. A negative control containing no *C. trachomatis* DNA was included. The PCR cycle conditions were as follows: 50°C for 2 min, 95°C for 5 min, 95°C for 3 s, and 60°C for 30 s. The last two steps were repeated for 40 cycles with fluorescence levels detected at the end of each cycle. Specificity of the primers was ensured with gel electrophoresis and with melting curve analysis. A standard curve was taken from purified *C. trachomatis* L2/434/Bu genomic DNA with serial dilutions for each gene-specific primer pair. The transcripts per genome copy were then calculated as the number of transcripts divided by the number of chlamydial genome copies measured with the same primer pair.

### Immunoblotting analysis

*C. trachomatis* infected cells in 24-well culture plate were lysed directly in 8 M urea buffer containing 10 mM Tris (pH 8.0), 0.1% SDS, and 2.5% β-mercaptoethanol. The protein content was determined by a bicinchoninic acid (BCA) protein assay kit (Thermal Fisher). The optimal amount of protein dissolved in 1× sodium dodecyl sulfate (SDS) loading buffer was separated on a 10% SDS-polyacrylamide gel and transferred to a polyvinylidene difluoride (PVDF) membrane (Millipore) for immunoblotting. The membrane was incubated with appropriate primary antibodies, followed by incubation with the secondary antibody that is conjugated with HPR. For complementation of *topA* knockdown, cells seeded in a 6-well plate were infected with *C. trachomatis* L2/topA-kdcom at an MOI of 1 in the presence of 500μg/mL spectinomycin and 1μg/mL cycloheximide. At 10h pi, cells were induced or not with 2nM (4 ng/mL) aTC. At 24h pi, protein lysates were harvested in 8M urea buffer with nuclease added immediately before use. Protein concentrations were quantified using EZQ protein assay kit (ThermoFisher) according to the manufacturer’s instructions. A total of 30μg protein per sample were separated on a 12% SDS-PAGE gel and then transferred to a PVDF membrane. The protein of interest was probed using goat anti-MOMP and rabbit anti-His_6_ antibodies followed by donkey anti-goat 680 and donkey anti-rabbit 800 secondary antibodies (LICOR, Lincoln, NE). The blot was imaged on an Azure c600 imaging system.

### Statistical analysis

Data for the assays include the mean ± standard derivation of at least two independent experiments. For multiples comparisons, one-way analyses of variance (ANOVA) with 95% significance level were performed. GraphPad Prism was used for all analyses. Differences were considered statistically significant when *P* <0.05.

## ACKNOWLEDGMENTS

Research reported in this publication was supported in part by National Institutes of Allergy and Infectious Diseases grants AI146454 to LS and a National Science Foundation CAREER grant (1810599) to SPO.

