## supplemental Table S1-S2 and Supplemental Figure S1-S5 for "Targeted repression of DNA topoisomerase I by CRISPRi reveals a critical function for it in the *Chlamydia trachomatis* developmental cycle"

**Table S1 and Table S2**

**Figures S1-S5**

**Table S1. Strains and plasmids used in this study**

| Strains | Relevant Details | Reference |
|---|---|---|
| ***C. trachomatis*** | | |
| L2/434/Bu | Bubo isolate from human with lymphogranuloma venereum (LGV). | ATCC VR-902B™ |
| L2/topA-kd | Transformed LGV/L2 with a CRISPRi plasmid encoding *topA*-specific crRNA and $P_{tet}$-directed synthesis of dCas12 protein | This study |
| L2/topA-Nt | Transformed LGV/L2 as a vector control | This study |
| L2/topA-kdcom | Transformed LGV/L2 with a plasmid encoding *topA-his6* for complementation | This study |
| ***E. coli*** | | |
| 10-beta (Catalog C3019H) | Host cell for cloning (*araD139D(ara-leu)7697fhuA lacX74 galK (f80 D(lacZ)M15) mcrA galU recA1 endA1 nupG rpsL (*$Str^r$*) D(mrr-hsdRMS-mcrBC)* | New England Biolabs |
| **Plasmid** | | |
| pBOMBL12CRia(e.v.)::L2 (aka pBOMBL-As_dCas12::L2) | *aadA* $P_{tet}$-directed synthesis of dCas12 protein; $P_{Nmen}$::*gfp* ($Spc^r$) | This study |
| pBOMBL12CRia(NT)::L2 | *aadA* $P_{Nmen}$::*gfp* $P_{tet}$::As_dCas12vaa $P_{dnaKmut}$::As_crRNA_non-targeting ($Spc^r$) | This study |
| pBOMBL12CRia(topA)::L2 | *aadA* $P_{Nmen}$::*gfp* $P_{tet}$::As_dCas12vaa $P_{dnaKmut}$::As_crRNA_topA ($Spc^r$) | This study |
| pBOMBL12CRia-topA_6xH(topA)::L2 | *aadA* $P_{Nmen}$::*gfp* $P_{tet}$::As_dCas12vaa-topA_6xH $P_{dnaKmut}$::As_crRNA_topA ($Spc^r$) | This study |

### Table S2. Primers and gBlocks used in this study

| Primer name | sequence 5'→3' | Use/features |
|---|---|---|
| euo-rtF | TCAAGGAGAGCTTCTGTTTGATAAC | RT-qPCR for *euo* |
| euo-rtR | TGCGTGTAGCATAGTAAATCTTCTG | |
| tuf-rtF | GTAACTCTGCCTGAGGGAATTGA | RT-qPCR for *tufA* |
| tuf-rtR | CACGAATCGCAAATCTCATACCT | |
| gfp_rtF | GTATACATCATGGCAGACAAACAA | RT-qPCR for *gfp* |
| gfp_rtR | TGTTGATAATGGTCTGCTAGTTGAA | |
| incD_rtF | CTCTGTAGCCCTGTTTCTGTTTTAG | RT-qPCR for *incD* |
| incD_rtR | CTAGTCACAGCTTCTGTAGTCAGCA | |
| omcB_rtF | GTTTGCGTTGCCCAGTAGTT | RT-qPCR for *omcB* |
| omcB_rtR | CACGCTGTCCAGAAGAATGA | |
| hctB_qRT_F | AAACATACTGCAGCTTGTGGAC | RT-qPCR for *hctB* |
| hctB_qRT_R | GAGCTGTACGAGAACGGTTAGG | |
| rtct190/189PrF | GAGTCACGCTTTATCCATTCGG | RT-qPCR for *gyrB/gyrA* |
| rtct190/189PrR | AGCTCTCCTTCATTTCCTCTTCA | |
| rtCT643prF | GTTGAATCCCCAGCCAAGATTA | RT-qPCR for *topA* and sequencing |
| rtCT643prR | CCCTTTTGCAGGAAGATCAACA | |
| rtCT660/661PrF | GAGAATCTTGTTACCAACCTCTAGC | RT-qPCR for *pare/parC* |
| rtCT660/661PrR | GTTCCAAAATGACGTAAGACGC | |
| topA_rtF2 | CAATCGCATACCAAGCCTTT | RT-qPCR for *topA* and sequencing |
| topA rtR2 | ACGTAGCCTTTGCCCTTCTTT C | |
| topA/(dCas12vaa)/5' | cgcaacgtagctgcttaagtaccgaggagaatctgcATGAAAAAATCCTTAATCATTG | Cloning *topA-his6* |
| topA_6xH/(pL12CRia)/3' | catgagcggatacatatttgaatggttaatggtgatggtgatggtgCTCTTCCTTGATTAAGTGCG | |

| gBlock Name | sequence 5'→3' | Use/Features |
|---|---|---|
| *topA* crRNA | tgtgaaagtgggtcttaagacgtcggtactgcatgtgacgcacgtagatcatgca*TTCACCGGTGGAGACGGTTTTCTTATAATGACACC*TAATTTCTACTCTTGTAGAT**TGCGAGAGACTAAGATCCCGCC**CAAATAAAACGAAAGGCTCAGTCGAAAGACTGGGCCTTTCGTTTTATcaacagcggtctactgaatctgagctagtgcgtgatataattaaaattatattca | For CRISPRi knockdown of *topA;*<br><br>Lower case for plasmid overlap and spacer, *italicized* for $P_{dnaK}$ sequence, underlined for crRNA scaffold, **bold** for *topA* targeting sequence, Upper case for *rrnB*1 terminator |
| Non-targeting crRNA | tgtgaaagtgggtcttaagacgtcggtactgcatgtgacgcacgtagatcatgca*TTCACCGGTGGAGACGGTTTTCTTATAATGACACC*TAATTTCTACTCTTGTAGAT**ACCGAGTTGCCCGTTAAAGTA**CAAATAAAACGAAAGGCTCAGTCGAAAGACTGGGCCTTTCGTTTTATcaacagcggtctactgaatctgagctagtgcgtgatataattaaaattatattca | Non-targeting CRISPRi control;<br><br>Lower case for plasmid overlap and spacer, *italicized* for $P_{dnaK}$ sequence, underlined for crRNA scaffold, **bold** for *non-targeting* targeting sequence, Upper case for *rrnB*1 terminator |

**Supplemental figures:**

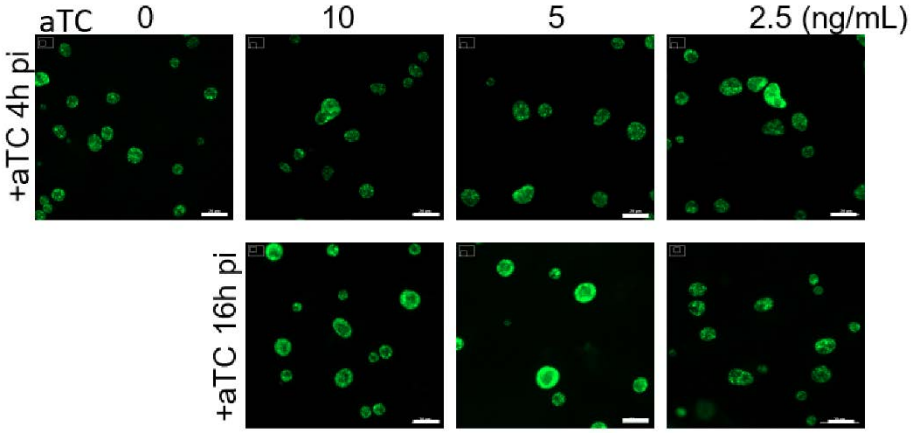


**Figure S1.** Live-cell images of *C. trachomatis* L2/Nt that has wild-type *topA* expression. HeLa cells were infected with *C. trachomatis* L2/Nt at MOI ~0.4 and cultured in aTC free medium. The increasing concentrations of aTC (0, 2.5, 5, or 10ng/mL) were added starting at 4 h pi (upper panels) or 16 h pi (lower panels). The automated imaging was taken at 24 h pi under the same exposure conditions using Cytation 1. Scale bar=20µm

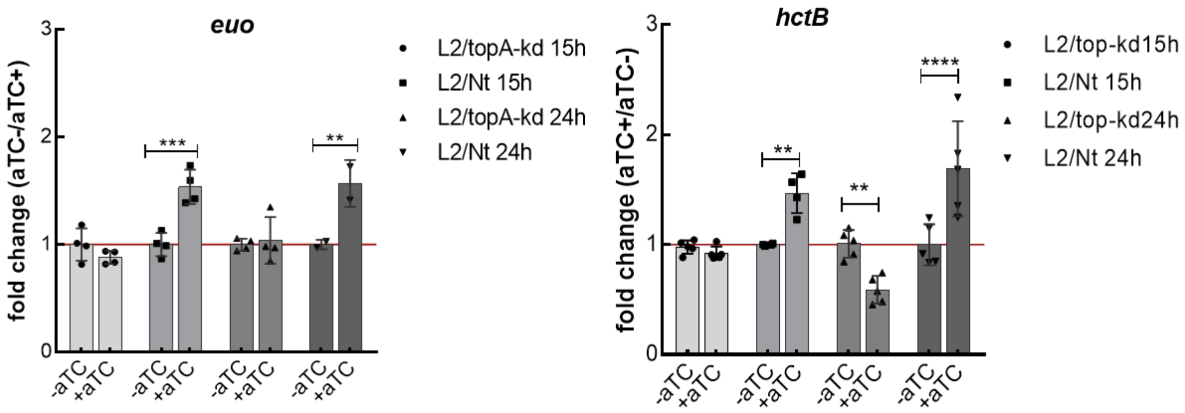


**Figure S2**. Quantifying transcript levels of *euo* and *hctB* in *C. trachomatis* using RT-qPCR. The mRNA transcript levels were normalized to the DNA control as determined by qPCR targeting chlamydial *tufA*. Values were presented as mean ± SD of four biological replicates. Statistical significance in all panels was determined by one-way ANOVA followed by Tukey's post-hoc test. **p < 0.01, ***p < 0.001, ****p < 0.0001.

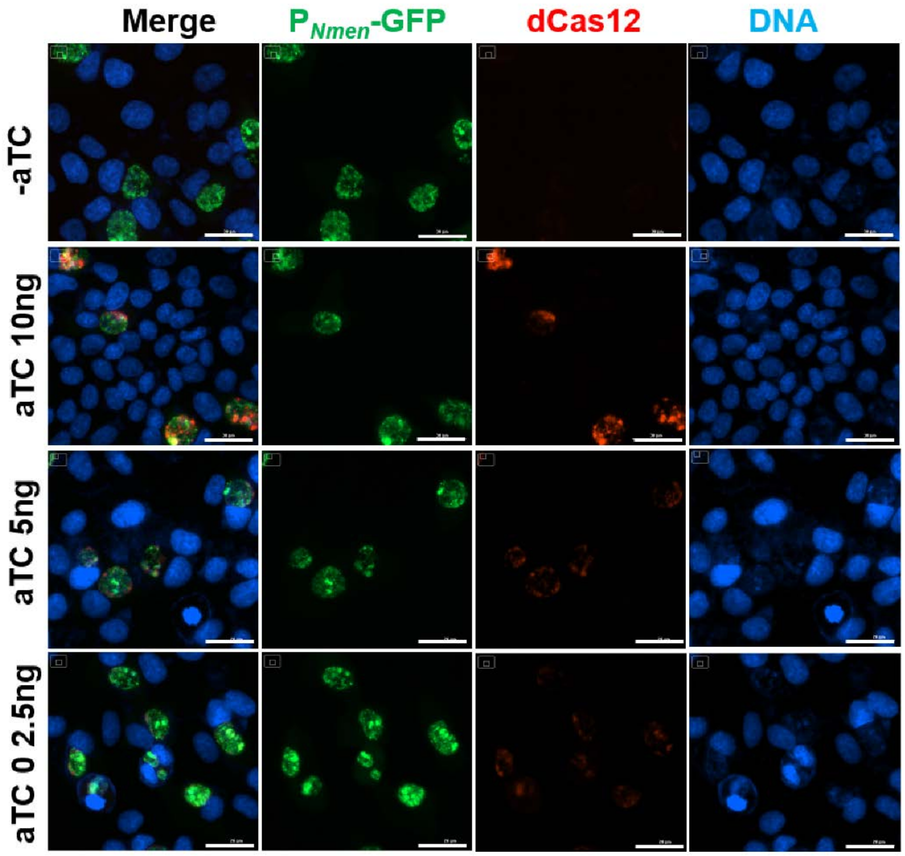


**Figure S3**. Immunofluorescence micrographs of *C. trachomatis* L2/topA-kdcom expressing dCas12. Cells were grown in the absence or presence of increasing concentrations of aTC and fixed at 40 h pi for IFA. The dCas12 was immunolabelled with rabbit anti-dCas12 antibody and visualized with Alexa Fluor 568-conjugated goat anti-rabbit IgG. DAPI-stained DNA (blue), *C. trachomatis* expressing GFP (green), and dCas12 (red) are shown. Scale bar=30μm.

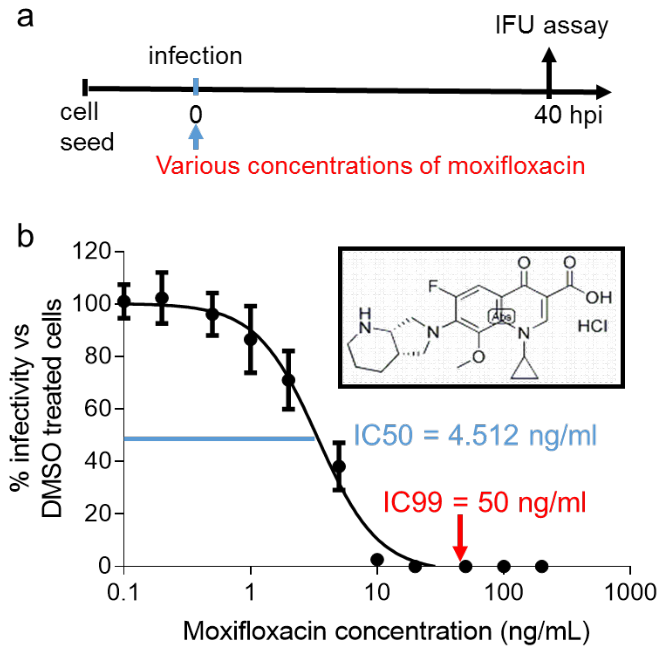


**Figure S4.** Determining MICs of moxifloxacin against *C. trachomatis.* (**a**) Schematic diagram of the experimental procedure. Moxifloxacin was added into the *C. trachomatis* culture immediately after infection. (**b**) MIC of moxifloxacin. Structure of moxifloxacin is shown. *C. trachomatis* infected HeLa cells were cultured in medium containing moxifloxacin (at the concentrations of 0.1, 0.25, 0.5, 1, 2.5, 10, 25, 50, 100, and 400 ng/mL) for 40 h prior to the analysis of IFUs by passaging on a fresh monolayer of HeLa cells in the absence of antibiotic. The relative infectivity (x-axis) normalized to DMSO control was presented as percentage (mean ± SD). The moxifloxacin concentrations are shown on the y-axis. Studies were repeated three times in quadruplicate.

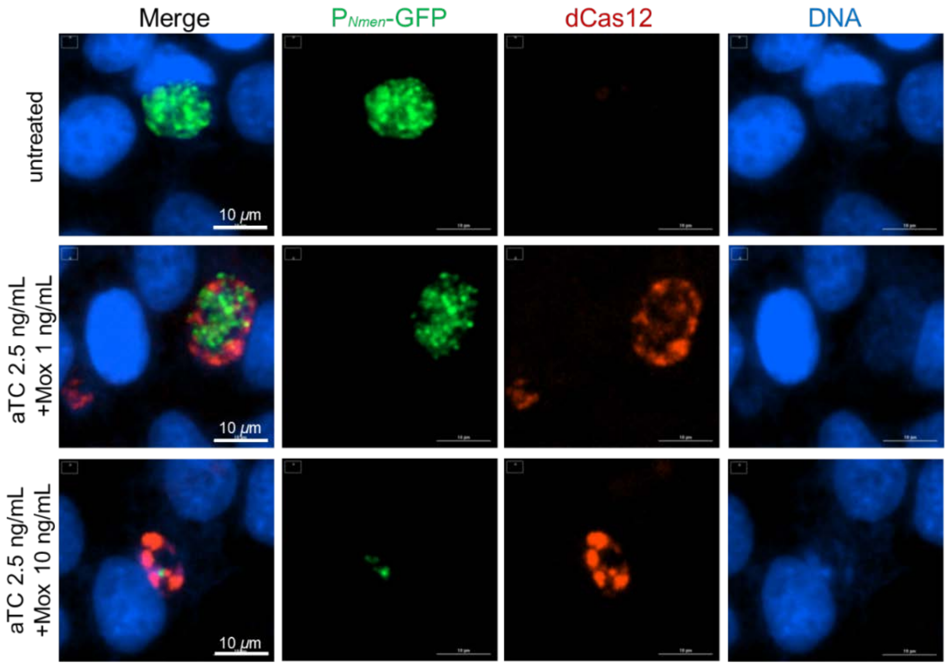


**Figure S5**. Immunofluorescence micrograph of *C. trachomatis* L2/topA-kdcom expressing dCas12. HeLa cells with infection of *C. trachomatis* were cultured in the presence of aTC (at 2.5 ng/mL) and Mox (at 1 ng/mL or 10 ng/mL) for 40 h pi and fixed for IFA. The dCas12 was immunolabelled with rabbit anti-dCas12 antibody and visualized with Alexa Fluor 568-conjugated goat anti-rabbit IgG. DAPI-stained DNA (blue), *C. trachomatis* expressing GFP (green), and dCas12 (red) are shown. Scale bar=10µm.
